# Setdb1 safeguards genome integrity in muscle stem cells to allow for regenerative myogenesis and inflammation

**DOI:** 10.1101/2023.06.08.544190

**Authors:** Pauline Garcia, William Jarassier, Caroline Brun, Lorenzo Giordani, Fany Agostini, Wai Hing Kung, Cécile Peccate, Jade Ravent, Sidy Fall, Valentin Petit, Tom H Cheung, Slimane Ait-Si-Ali, Fabien Le Grand

## Abstract

Modulations in chromatin structure orchestrate gene expression and direct stem cell fate. More specifically, the histone 3 lysine 9 Methyltransferase Setdb1 controls transcriptional repression to regulate pluripotency and self-renewal. While Setdb1 functions have been extensively studied in embryonic stem cells and in cancer cells, less is known on its role in adult stem cells *in vivo*. Here, we show that Setdb1 expression by adult muscle stem cells (MuSCs) is required for muscle tissue regeneration following acute injury. We find that SETDB1 represses the expression of the endogenous retroviruses (ERVs) family of transposable elements in MuSCs. ERV de-expression in *Setdb1-null* MuSCs prevents their amplification following exit from quiescence and promotes cell death. Multi-omics profiling further shows that the absence of SETDB1 in MuSCs leads to the activation of the DNA-sensing cGAS-STING pathway, entailing increased cytokine expression. *In vivo*, conditional disruption of *Setdb1* in MuSCs provokes aberrant infiltration of inflammatory cells including the appearance of a pathological macrophage lineage. The ensuing histiocytosis is accompanied by necrosis of the newly formed muscle fibers which, in addition with the progressive loss of MuSCs, completely abolish skeletal muscle tissue repair. In contrast, disruption of *Setdb1* gene in another muscle-resident cell type, the fibro-adipogenic progenitors (FAPs), does not impact regenerative inflammation. In conclusion, the control of genome stability by SETDB1 in an adult somatic stem cell is necessary for both its regenerative potential and an adequate inflammation regulating tissue repair.

## INTRODUCTION

The timing of gene expression in stem cells is crucial for proper lineage commitment, development and regeneration.^1^ Chromatin structure regulates gene expression and stem cell fate choices. As such, chromatin deregulation is linked to developmental defects, diseases, and aging.^2^

Lysine methyltransferases (KMT) and demethylases (KDM) are critical regulators of chromatin. Histone 3 methylation at lysine position 9 (H3K9) is necessary for the assembly of constitutive and facultative heterochromatin. SUV39 family KMTs, including SUV39H1/2, G9A, GLP, and SETDB1/2, primarily methylate H3K9. Although these KMTs can substitute for each other in some cases, their functions are not interchangeable. SETDB1 (SET domain bifurcated histone lysine methyltransferase 1, also called ESET or KMT1E) is responsible for H3K9 di- and tri-methylation involved in gene repression in euchromatin and facultative heterochromatin.^3^ SETDB1 is required for transcriptional repression, pluripotency, and self-renewal of mouse embryonic stem cells (mESCs).^4^ Gain-of-function mutations of *Setdb1* are associated with various types of cancer, promoting cell cycle, proliferation, migration, and invasion of cancer cells.^5,6^ Yet, little is known about Setdb1 roles in adult stem cells *in vivo*, and no studies have investigated its impact on adult tissue regeneration following injury.

Adult skeletal muscle consists of multinucleated innervated myofibers that ensure muscle functions and contractility. Myofibers host a population of quiescent muscle stem cells (MuSCs) in a niche between their plasma membrane and the surrounding basal lamina.^7^ MuSCs have the ability to repair and form *de novo* myofibers to regenerate damaged muscle tissue. Following injury of their host myofibers, MuSCs activate from quiescence, give rise to transient amplifying muscle progenitor cells (also called myoblasts) that will differentiate and fuse into myotubes that subsequently mature into myofibers.^8^ Understanding the molecular and cellular mechanisms that regulate MuSC activation, proliferation, and differentiation is crucial for developing therapies against neuro-muscular diseases.

Previous research found that SETDB1 is highly expressed in activated MuSCs, primarily nuclear, but relocates to the cytoplasm upon myoblast differentiation in a canonical WNT signaling-dependent manner.^9^ This differential localization of SETDB1 corresponds to a change in the gene expression program required for myoblast terminal differentiation. However, the role of SETDB1 in proliferative MuSCs during muscle regeneration is unknown. To investigate it, we generated a conditional mouse model that allows the specific disruption of *Setdb1* in MuSCs.

## RESULTS

### SETDB1 is required for muscle regeneration

To study the impact of *Setdb1* gene silencing in MuSCs during skeletal muscle regeneration, we crossed mice harboring a floxed *Setdb1* allele^10^ with mice expressing CreERT2 from the *Pax7* locus as a bicistronic transcript^11^ (Figure S1A). To induce recombination at the *Setdb1* loci in MuSCs, we injected 10-week-old *Pax7*^CreERT2/+:^*Setdb1*^lox/lox^ mice with tamoxifen (*Setdb1^MuSC-KO^*) diluted in corn oil. For controls, we injected mice of the same genotype with corn oil only (*Control*). Genotyping PCR of MuSCs isolated from tamoxifen-induced mice showed an 85% deletion of *Setdb1* allele as compared to oil-treated mice (Figure S1B), which is efficient but not complete. As binding partners of SETDB1 are involved in neuronal progenitors homeostasis^12^, we first analyzed the neuro-muscular junctions (NMJ) of myofibers. We did not see any difference in NMJ compactness nor fragmentation index between *Setdb1^MuSC-KO^* and *Control* myofibers. We hence conclude that the loss of *Setdb1* in MuSCs does not affect NMJ (Figure S1C).

MuSCs are mostly quiescent, as only a minute proportion (2 to 4%) can be seen cycling in adult muscles.^13^ As such, we examined if conditional *Setdb1* gene disruption could induce MuSCs to exit quiescence (Figure S1D, S1E). Three months after tamoxifen treatment, *Control* and *Setdb1^MuSC-KO^* mice exhibited similar numbers of sublaminar PAX7+ MuSCs (Figure S1F). Furthermore, we did not observe any alterations of anatomy, histology and myofiber size in *Setdb1^MuSC-KO^ Tibialis anterior* (TA) muscles (Figure S1E, S1G). Together, our observations indicate that *Setdb1* gene disruption does not perturb the quiescent MuSC pool and therefore, SETDB1 is not required to control MuSC quiescence.

To determine if *Setdb1^MuSC-KO^* can support injury-induced adult regenerative myogenesis, we injured TA muscles using cardiotoxin^14^ and sampled the tissues 28 days post-injury (d.p.i.) (Figure 1A). While *Control* muscles appeared well regenerated, *Setdb1^MuSC-KO^* muscles were severely reduced in size (Figure 1B), exhibiting more than a 50% decreased mass compared to either the contralateral muscle or to the regenerated *Control* muscle (Figure 1C).

**Figure 1.**
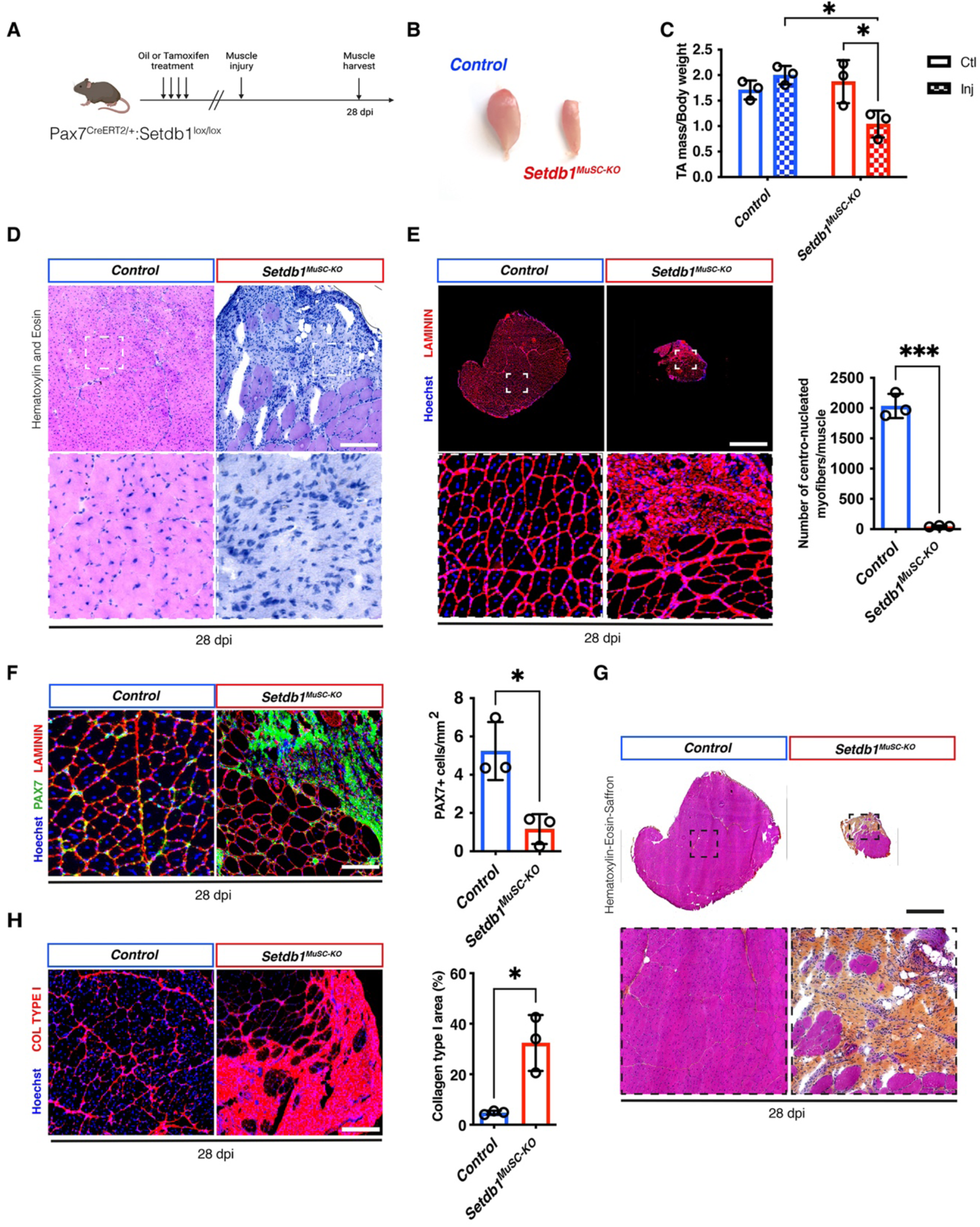
The loss of SETDB1 in MuSC leads to the abolition of muscle regeneration. A. Experimental scheme; *Pax7^CRE-ERT2^; Setdb1^fl/fl^* mice intraperitoneally injected with either Oil or Tamoxifen and cardiotoxin (CTX) is performed on TA (*Tibialis Anterior*) muscle after 28 days post injury. B. Representative photographs representing TA muscle after 28 days post injury in both conditions. C. Normalized TA weight on mice body weight represented in *Control* and *Setdb1^MuSC-KO^* conditions in uninjured and injured TA (n=3 mice). D. Hematoxylin and eosin stain of TA muscle cross sections at 28 days post injury (CTX) (n=3 mice). E. Immunostaining for LAMININ on TA cryosections 28 days after injury. Quantification of the number of centro-nucleated myofibers on TA cryosections 28 days after injury (n=3 mice). F. Quantification of PAX7+ cells on TA cryosections for the *Control* and *Setdb1^MuSC-KO^* conditions (n=3 mice). G. Hematoxylin-Eosin-Saffron stain of TA cryosections 28 days after injury (n=3 mice). H. COLLAGEN I immunostaining on TA cryosections 28 days after injury (n=3 mice) and quantification of the COLLAGEN I area in *Control* and *Setdb1^MuSC-KO^* conditions. Data are represented as mean ± SD; *p < 0.05, ***p < 0.001, t test. Scale bars, (D) 150 μm, (E) 200 μm, (F) 100 μm, (G) 200μm, (H) 150 μm.

Histology of tissue cross-sections showed that deletion of *Setdb1* in MuSCs completely abrogated the tissue regeneration process (Figure 1D, S1H). As such, injured *Setdb1^MuSC-KO^* muscles did not contain any new myofibers as compared to injured controls that were mostly composed of regenerated myofibers with centrally-located nuclei (Figure 1E). This impairment of the regenerative process was accompanied by a 5-fold reduction in the numbers of PAX7+ MuSCs (Figure 1F). Investigation of the nature of the residual tissue composing the injured *Setdb1^MuSC-KO^* muscles with Hematoxylin-Eosin-Saffron staining revealed that myofibers were replaced by a scar composed of connective tissue (Figure 1G). As such, quantification of Collagen Type I immunoreactive area showed fibrosis of injured *Setdb1^MuSC-KO^* muscles (Figure 1H). Thus, *Setdb1* expression in MuSCs is absolutely required for injury-induced adult myogenesis.

### Loss of *Setdb1* expression in MuSCs delays myofiber formation

To evaluate at which stage the regeneration is nullified in *Setdb1^MuSC-KO^* mice, we performed injury experiments and sampled the regenerating tissues at early time-points (Figure 2A). In these experimental conditions, we validated that *Setdb1* expression is downregulated *in vivo* in MuSCs isolated by fluorescence-activated cell sorting (FACS) from regenerating tissues (Figure S2A).

**Figure 2.**
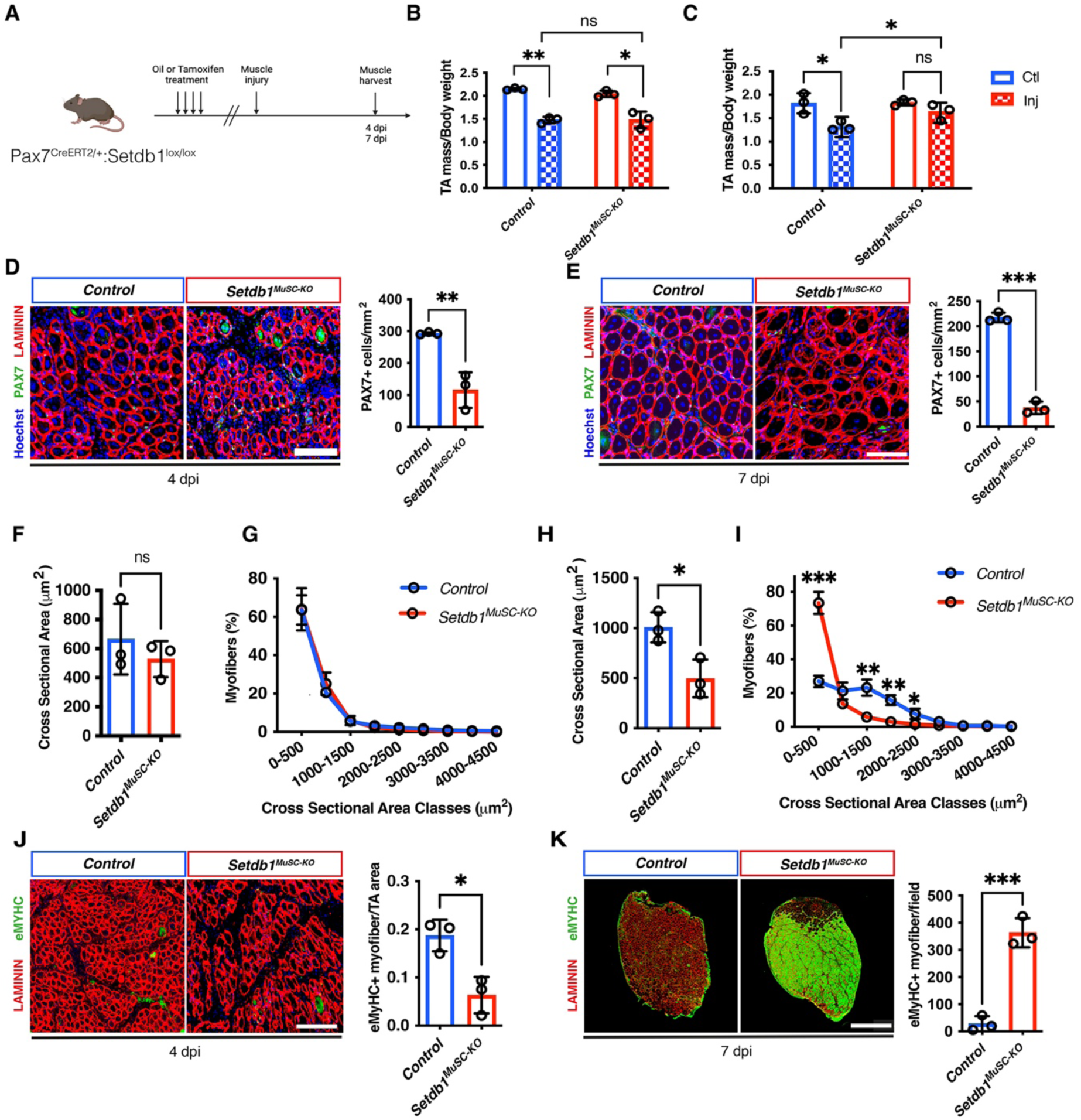
*Setdb1* gene disruption in MuSCs induces cell death and muscle regeneration delay. A. Experimental set up; *Pax7^CRE-ERT2^; Setdb1^fl/fl^* mice intraperitoneally injected with either Oil or Tamoxifen. Cardiotoxin-induced injury (CTX) was performed in the TA (*Tibialis Anterior*) muscles and analyzed 4- and 7-days post-injury. B. Normalized TA weight on mice body weight represented in *Control* and *Setdb1^MuSC-KO^* muscles in uninjured and injured TA after 4 days post injury (n=3 mice). C. Normalized TA weight on mice body weight represented in *Control* and *Setdb1^MuSC-KO^* muscles in uninjured and injured TA after 7 days post injury (n=3 mice). D. LAMININ and PAX7 immunostaining in *Control* and *Setdb1^MuSC-KO^* muscles after 4 days of the CTX injury and quantification of the PAX7+ cells in both conditions (n=3 mice). E. LAMININ and PAX7 immunostaining in *Control* and *Setdb1^MuSC-KO^* muscles after 7 days of the CTX injury and quantification of the PAX7+ cells in both conditions (n=3 mice). F. Cross sectional area of CTX-injured TA muscles of *Control* and *Setdb1^MuSC-KO^* muscles after 4 days post injury (n=3 mice). G. Distribution of cross-sectional area of CTX-injured TA muscles, as determined in F (n=3 mice). H. Cross sectional area of CTX-injured TA muscles of *Control* and *Setdb1^MuSC-KO^* muscles after 7 days post injury (n=3 mice). I. Distribution of cross-sectional area of CTX-injured TA muscles, as determined in F (n=3 mice). J. eMYHC and LAMININ immunostaining in *Control* and *Setdb1^MuSC-KO^* muscles at 4 days after injury and quantification of the number of eMYHC+ myofibers per cryosections (n=3 mice). K. eMYHC and LAMININ immunostaining in *Control* and *Setdb1^MuSC-KO^* muscles at 7 days after injury and quantification of the number of eMYHC+ myofibers per field (n=3 mice). Data are represented as mean ± SD; ns not significant, *p < 0.05, **p < 0.01, ***p < 0.001, t test. Scale bars, (D) 100 μm (E) 20 μm (J) 50 μm (K) 400 μm.

Intriguingly, we did not observe a difference in the mass of *Setdb1^MuSC-KO^* muscles compared to *Control* muscles at 4 d.p.i. (Figure 2B). At 7 d.p.i., the weights of injured *Setdb1^MuSC-KO^* muscles were closer to their contralaterals than to *Control* ones (Figure 2C). However, examination of regenerating tissues cross-sections revealed perturbations in the regeneration process in *Setdb1^MuSC-KO^* muscles. Indeed, we observed a drastic loss of MuSCs within *Setdb1^MuSC-KO^* muscles as compared to *Controls* at both 4 and 7 d.p.i. (Figure 2D, 2E). Evaluation of the cell cycle state of PAX7+ cells on tissue sections showed reduced MuSC proliferation in *Setdb1^MuSC-KO^* muscles (Figure S2C). We found the regenerating *Setdb1^MuSC-KO^* muscles to be disorganized compared to *Control* muscles at 4d.p.i (Figure 2D, S2B) with myofibers displaying aberrant shapes in *Setdb1^MuSC-KO^* muscles at 7d.p.i. (Figure 2E, Figure S2D). This disorganization was accompanied by increased amounts of PDGFRα+ fibro-adipogenic progenitors (FAPs) (Figure S2E) and of fibrotic tissue (Figure S2F) in regenerating *Setdb1^MuSC-KO^* muscles compared to *Control* muscles. In contrast, while measurement of the newly-formed myofibers size did not show any difference between the different conditions at 4 d.p.i. (Figure 2F, 2G, S2B), we observed severe atrophy of regenerated *Setdb1^MuSC-KO^* myofibers at 7.d.p.i. (Figure 2H, 2I, S2D). From these results, we conclude that muscle regeneration is strongly affected in the absence of Setdb1 expression by MuSCs, although atrophied new myofibers can still be formed.

To evaluate the maturation of regenerated myofibers, we performed immunostaining against the embryonic Myosin Heavy Chains (eMyHC). This Myosin isoform is specifically expressed by newly formed myofibers during regeneration at early timepoints and is absent from well-regenerated myofibers. At 4 d.p.i., a strong reduction of the number of eMyHC+ myofibers was found in the *Setdb1^MuSC-KO^* muscles compared to *Control* muscles, indicating a delay in myogenesis (Figure 2J). Surprisingly, at 7 d.p.i., almost all of the regenerating myofibers were positive for eMyHC+ in *Setdb1^MuSC-KO^* muscles, while it had become hardly detectable in *Control* muscles (Figure 2K). The relative expression levels of *Pax7*, *Myh4* (adult isoform), *Myh8* (peri-natal isoform) obtained by quantitative RT-PCR (qRT-PCR) on cDNA prepared from muscle samples at 7 d.p.i are in accordance with the immunolocalization results (Figure S2G).

Taken together, these results show that MuSC-specific *Setdb1* gene disruption leads to impaired muscle regeneration accompanied by fibrosis. However, *de novo* myofiber formation is still present, albeit severely perturbed.

### MuSC proliferation and viability rely on SETDB1

To understand the consequences of *Setdb1* gene disruption in MuSCs, we sorted MuSCs from *Control* and *Setdb1^MuSC-KO^* muscles using magnetic-activated cell sorting (MACS) (Figure 3A). While we observed no difference at day 3 post-extraction, *Setdb1^MuSC-KO^* MuSCs presented a striking deficit in colony formation *in vitro* after 7 days (Figure 3B, Figure S3A). This almost complete absence of *in vitro* amplification potential precluded more detailed analysis of *Setdb1^MuSC-KO^* MuSCs phenotype. To overcome this pitfall, we extracted MuSCs from untreated *Pax7*^CreERT2/+:^*Setdb1*^lox/lox^ mice and cultured them as primary myoblasts. Following *in vitro* amplification, myoblasts were then treated with either vehicle (EtOH) or 4-hydroxytamoxifen (4-OHT) for 3 days to induce *Setdb1* gene recombination *in vitro* (Figure 3C). 4-OHT treatment reduced the proliferation of primary cells *in vitro*, as shown by decreased EdU incorporation (Figure 3D, 3E) and decreased proportion of Ser10-phospho-histone H3+ cells (Figure S3B). Accordingly, qRT-PCR revealed that several cyclins (*Ccna1*, *Ccnb1*, *Ccnd1* and *Cdc25a*) expression levels were significantly reduced in *Setdb1^MuSC-KO^* cells while cyclin-dependent-kinase inhibitors (*Cdkn1a*, *Cdkn2a* and *Cdkn1b*) expression levels were increased (Figure 3F). Cell cycle analysis delineated a specific decrease in the fraction of cells in S-phase in *Setdb1^MuSC-KO^* cells compared to *Control* cells (Figure 3G). Importantly, immunostaining for PAX7 as well as gene expression analysis for *Pax7*, *Myod1* and *Myog* did not show any significant difference between vehicle*-*treated and 4-OHT-treated cells (Figure 3H-3I, Figure S3E). This indicates that the proliferation defect induced by *Setdb1* gene disruption is not related to a loss of myogenic identity. To evaluate if loss of *Setdb1* may render cells senescent, we tested for SA-b-gal activity in *Control* and *Setdb1^MuSC-KO^* cells but did not observe any difference between conditions (Figure S3C-S3D). We next stained cells with PI and YO-PRO-1 to evaluate the percentage of early and late apoptotic cells, as well as the number of necrotic cells, by flow cytometry. Interestingly, the *Setdb1^MuSC-KO^* cells exhibited an increase of the three conditions, demonstrating that the loss of *Setdb1* leads to cell death (Figure 3J-L).

**Figure 3.**
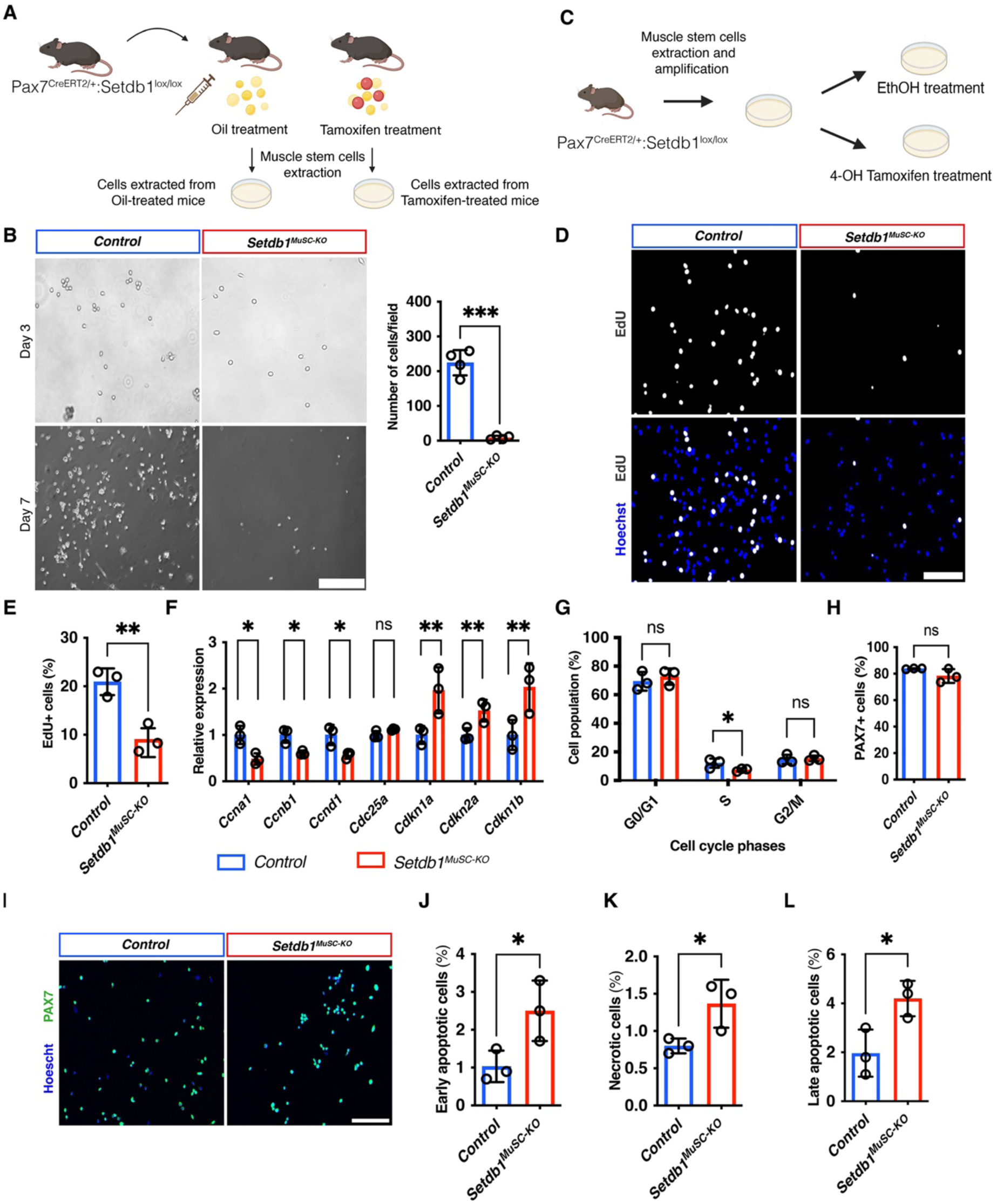
MuSC proliferation and viability are affected in absence of SETDB1. A. Experimental scheme of the set up used to extract myoblasts in culture *in vitro* after oil and Tamoxifen treatment of the *Pax7^CRE-ERT2^; Setdb1^fl/fl^* mice model. B. Photographs representing the extracted myoblasts after 3 days and 7 days *in vitro* from the *Control* and *Setdb1^MuSC-KO^* mice. Quantification of the cell number in the *Control* and *Setdb1^MuSC-KO^* cells *in vitro* after 7 days (n=4). C. Experimental scheme used to analyze the myoblasts in culture *in vitro* in both conditions *Control* (treated with Ethanol (EtOH)) and *Setdb1^MuSC-KO^* (treated with 4-Hydroxitamoxifen (4-OHT)) condition. D. Immunostaining for EdU on myoblasts *in vitro* in both conditions *Control* (treated with Ethanol (EtOH)) and *Setdb1^MuSC-KO^* (treated with 4-Hydroxitamoxifen (4-OHT)) condition. E. Quantification of the number of EdU+ cells *Control* and *Setdb1^MuSC-KO^* after EtOH and 4-OHT treatment (n=3). F. Relative expression of genes that positively (*Ccna1*, *Ccnb1*, *Ccnd1* and *Cdc25a)* or negatively (*Cdkn1a*, *Cdkn2a* and *Cdkn1b*) regulates cell cycle after 5 days of EtOH or 4-OHT treatment (n=3). G. Flow cytometry analysis of cell cycle phases obtained following Propidium iodide staining (n=3). H. Quantification of PAX7+ cells after 5 days of EtOH and 4-OHT treatment in the *Control* and *Setdb1^MuSC-KO^* cells (n=3). I. Immunostaining of PAX7+ cells after 5 days of EtOH and 4-OHT treatment (n=3). J. Flow cytometry plots using Propidium Iodide staining and YO-PRO and quantification of the early apoptotic cells in the *Control* and *Setdb1^MuSC-KO^* cells (n=3). K. Quantification of the late apoptotic cells in the *Control* and *Setdb1^MuSC-KO^* cells (n=3). L. Quantification of the necrotic cells in the *Control* and *Setdb1^MuSC-KO^* cells (n=3). Data are represented as mean ± SD; ns not significant, *p < 0.05, **p < 0.01, ***p < 0.001, t test. Scale bars, (B) 150 μm (D) 100 μm (I) 100 μm.

To check if *Setdb1* gene disruption could affect the myogenic differentiation process, we treated proliferating cells with either EtOH or 4-OHT for 3 days and then, shifted the cultures into differentiation medium for another 3 days. Of note, the starting number of cells was higher in the 4-OHT condition, to compensate for the reduced growth rate. Immunostaining for sarcomeric MyHC (Figure S3F), accompanied by the quantification of differentiated nuclei (Figure S3G) and of the numbers of fused myonuclei *per* myotubes (Figure S3H-S3I) did not reveal any difference between *Control* and *Setdb1^MuSC-KO^* conditions. We concluded that the loss of *Setdb1* did not impact myogenic differentiation. Taken together, our *ex vivo* and *in vitro* data highlight that Setdb1 is crucial for MuSC proliferation and viability.

### SETDB1 restricts chromatin accessibility at TE sites in MuSCs to prevent cGAS-STING activation

To understand the molecular basis of the observed phenotype, we performed RNA-sequencing (RNA-seq) on MuSCs isolated by FACS from 60 hours-regenerating muscles (Figure S4A, S4B). Of note, RNA-seq validated reduced *Setdb1* gene expression by *Setdb1^MuSC-KO^* cells *in vivo* (Figure S4C). We observed a moderate impact of *Setdb1* gene disruption on MuSC protein-coding gene expression, with 210 genes up-regulated and 29 down-regulated (Figure 4A). Correlating with the phenotype observed *in vitro* and *in vivo*, gene set analysis revealed enrichment for pathways related to “Apoptosis” and “p53” in *Setdb1^MuSC-KO^* cells (Figure 4B; Table 1). Likewise, *Setdb1^MuSC-KO^* cells showed reduced expression of genes associated with cell cycle progression (Figures S4D). *Setdb1^MuSC-KO^* cells also exhibited enrichment for pathways related to “Inflammation response”. We validated the increased expression of inflammatory cytokines coding genes by MuSCs using qRT-PCR (Figure S4E). Interestingly, *Setdb1^MuSC-KO^* cells showed increased “Xenobiotic metabolism” pathway (Figure 4B), associated with a massive upregulation of glutathione S-transferase genes (*Gsta4* and *Gstt1*) (Figure 4A). GSTs have a key role in the biotransformation of substances that are not naturally expected to be present within the organism, of which can be the products of transposable elements (TE)^15,16^. Since SETDB1 is critical for TE silencing in mESCs^10,17^, we analyzed the expression of repetitive elements. Remarkably, *Setdb1^MuSC-KO^* MuSCs displayed re-expression of TEs of the Long Terminal Repeat family (LTRs) (Figure S4F). Among these LTRs, we found transcripts of the endogenous retrovirus (ERVs) class, notably of the ERVK and ERV1 subfamilies (Figure S4E). The most statistically significant TEs re-expressed by *Setdb1^MuSC-KO^* MuSCs included elements already described as SETDB1 targets, such as *MuLV*, *RTLR6* or *MMERVK10C* (Figure 4C).^18,19^

**Figure 4.**
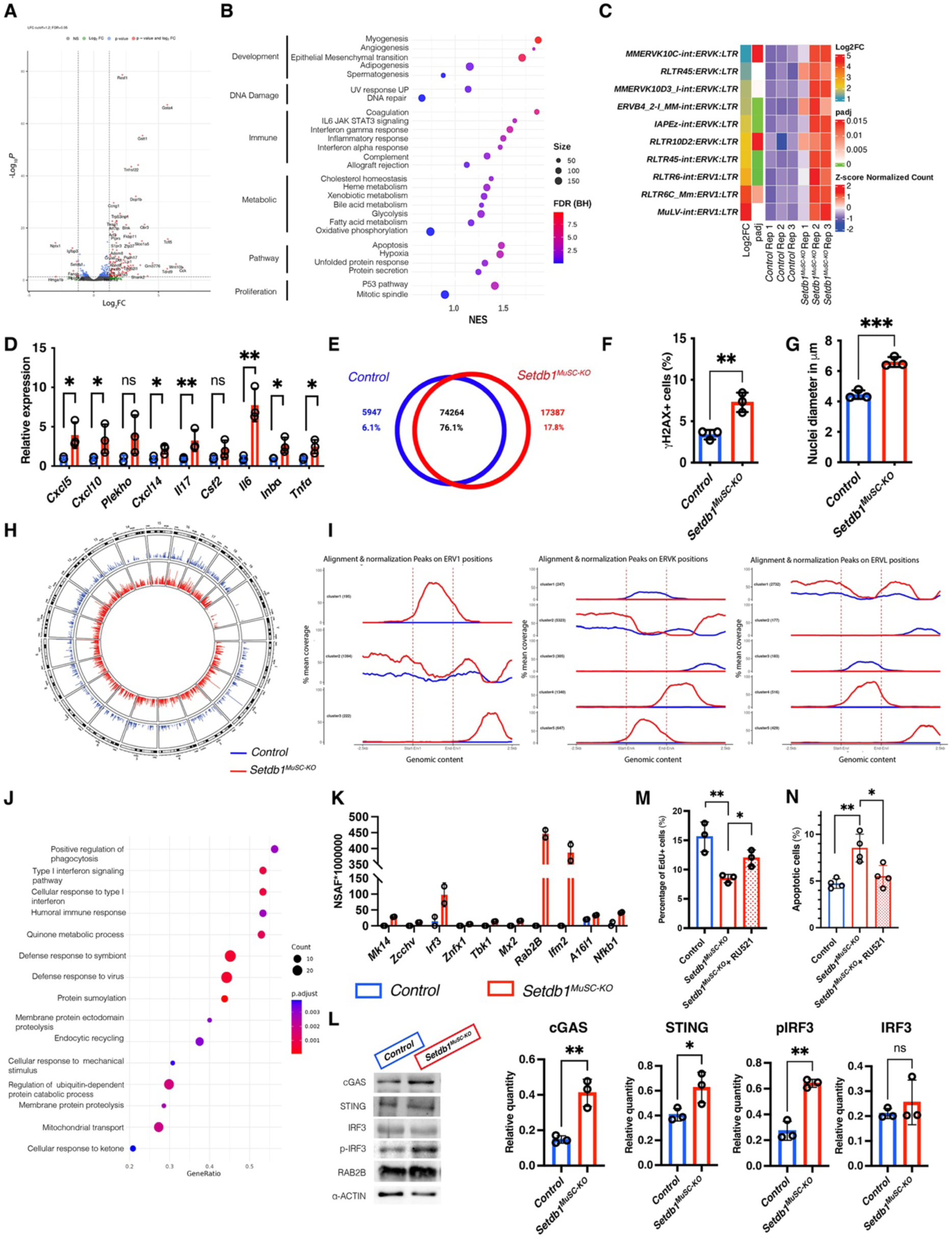
SETDB1 restricts chromatin accessibility at TE sites in MuSCs to prevent cGAS-STING activation. A. Volcano plot over *Setdb1^MuSC-KO^* muscles RNA-seq data (n=3). Dot in colors indicate significant dysregulated genes (FDR<0.05): in red, up-dysregulated (Log2 Fold-Change (LFC)>=1), in blue down-dysregulated (LFC<=-1). Dot in grey indicate no significant dysregulated genes or low value of LFC (abs(LFC)<1). B. Results of GSEA showing enriched gene sets from Mouse Hallmark on upregulated genes in *Setdb1^MuSC-KO^* muscles (n=3). Gene sets are grouped by process. Dot in red indicate a high significant enrichment (adjusted by FDR), dot in blue indicate a low significant enrichment. The size shows the number of gene dysregulated in each gene set and results are ordering in high to low level of Normalized Enrichment Score (NES). C. Heatmap of RNA-Seq normalized expression z-scores, log2FC and Pvalue computed for upregulated ERVs found in *Setdb1^MuSC-KO^* sorted MuSCs (n=3 mice). D. RT-qPCR of cytokines and interleukines found upregulated in the RNA-seq in *Control* and *Setdb1^MuSC-KO^* cells (n=3). E. Venn diagram showing overlapping ATAC-seq binding regions shared or not between *Control* and *Setdb1^MuSC-KO^* cells *in vitro.* F. Quantification of the γH2AX+ cells in *Control* and *Setdb1^MuSC-KO^* cells *in vitro* and in *Control* and *Setdb1^MuSC-KO^* cells (n=3). G. Measurement of the nuclei diameter in *Control* and *Setdb1^MuSC-KO^* cells *in vitro* (n=3). H. Genomic distribution of high ATAC-seq peaks (log10 (peak score) > 1.2) on mouse genome for *Control* and *Setdb1^MuSC-KO^* cells *in vitro* (n=2). I. ATAC signal coverage of the different ERV families found in the Control and *Setdb1^MuSC-KO^* cells (n=2). J. Results of GSEA showing enriched gene sets from Gene Ontology Biological Process on dysregulated protein in *Setdb1^MuSC-KO^* cells (n=2). Selected enriched terms are presented according to the fold enrichment. Dot in red indicate a high significant enrichment (adjusted by FDR), dot in blue indicate a low significant enrichment. The size shows the number of gene dysregulated in each gene set and results are ordering in high to low protein ratio observed/expected. K. Genes dysregulated in the *Setdb1^MuSC-KO^* cells in the “defense to virus” gene ontology analysis (n=2). L. Western blot analysis of cGas (60 kDa), STING (32 kDa), IRF3 (55 kDa), p-IRF3 (55kDa) and RAB2B (25kDa) in *Control* and *Setdb1^MuSC-KO^* cells *in vitro* and quantification in *Control* and *Setdb1^MuSC-KO^* cells *in vitro* (n=3). M. Percentage of EdU+ cells after 5 days of treatment with ethanol+DMSO, 4-OHT+DMSO or 4-OHT+RU521 at 200ug/ml *in vitro* (n=3). N. Percentage of apoptotic cells found in the Control, *Setdb1^MuSC-KO^* cells or *Setdb1^MuSC-KO^* cells treated with RU521 at 200ug/ml (n=4). Data are represented as mean ± SD; ns not significant, *p < 0.05, **p < 0.01, ***p < 0.001, t test.

As SETDB1 controls chromatin folding and subsequent gene repression, we investigated chromatin compaction at ERVs loci in *Control* and *Setdb1^MuSC-KO^*. To this end, we performed Assay for Transposase-Accessible Chromatin with sequencing (ATAC-seq) on cultured MuSCs treated with either EtOH or 4-OHT *in vitro* (Figure S4H, S4I). Of note, we chose not to perform ATAC-seq on FACS-sorted cells, since a high prevalence of dying cells is incompatible with ATAC-seq. Peak calling and differential analysis of accessible sites demonstrated a global increase in chromatin accessibility in *Setdb1^MuSC-KO^* MuSCs compared to *Control* cells (Figure 4D). Evaluating the signals specific to each condition showed *Setdb1^MuSC-KO^* cells displayed 17,387 specific peaks on the 97,598 total peaks, suggesting that the loss of *Setdb1* leads to chromatin opening at specific loci (Figure 4E; Table 2; Table 2bis). This release of chromatin compaction in *Setdb1^MuSC-KO^* cells was accompanied by increased DNA damage and enlarged nuclei diameter (Figure 4F, 4G). In accordance with gene expression results, visualization of enriched pathways from differentially accessible chromatin highlighted “Regulation of chemokine mediated signaling” in *Setdb1^MuSC-KO^* cells (Figure S4J).

We next visualized the distribution of peak annotations between samples, and found an increased proportion of distal intergenic regions in *Setdb1^MuSC-KO^*, where TE and ERVs are mostly located (Figure S4K). Through analyzing TEs specifically, we found around Transcription Start Site (TSS) localized in opened regions of the *Setdb1^MuSC-KO^*, TEs that belong to different ERV families (Figure 4L). Finally, we curated the TEs belonging to ERV1, ERVK and ERVL families into sub-clusters based on their ATAC signal coverage. We observed increased chromatin accessibility at specific ERV clusters in *Setdb1^MuSC-KO^* cells (Figure 4H). These results confirmed that the absence of SETDB1 modifies chromatin accessibility that induces the re-opening of ERV-containing regions.

We next asked if the increased chromatin accessibility is due to loss the heterochromatin mark H3K9me3 established by SETDB1. To test this, we profiled H3K9me3 marks using chromatin immunoprecipitation with sequencing (ChIP-seq). Interestingly, we observed a moderately lower (20%) peak number in the *Setdb1^MuSC-KO^* samples compared to *Control* samples (Figure S4M). Examination of the H3K9me3 peaks distribution along the genome indicated the heterochromatin marks were redistributed in *Setdb1^MuSC-KO^* cells (Figure S4O). In line with these results, *Setdb1^MuSC-KO^* cells did not show significant drop in the total abundance of H3K9me3 marks as assayed by immunolocalization (Figure S4N). We next calculated the overlap H3K9me3 coverage at TE loci, curated by sub-families. Interestingly, while H3K9me3 pattern did not change at non-ERV loci (such as LINE elements), we found the H3K9me3 marks to be less abundant at ERV loci (ERV1, ERVL) in *Setdb1^MuSC-KO^* cells (Figure S4P). These results suggest that de-repression of ERVs results from focal re-arrangement of H3k9me3 marks.

To understand the consequences of ERV de-repression in MuSCs, we performed mass-spectrometry-based proteomics on cultured MuSCs treated with either EtOH or 4-OHT *in vitro* (Figure S4Q, S4R). We identified 260 significantly deregulated proteins in *Setdb1^MuSC-KO^* samples compared to the *Control* samples (Figure 4J; S4S; Table 3). In line with the RNA-seq and ATAC-seq results, pathway analysis of the proteomics dataset also revealed enrichment for “Immune response” terms in *Setdb1^MuSC-KO^* cells (Figure 4I). Interestingly, the most significantly abundant and significant pathways were “Defense response to symbiont” and “Defense response to virus” (Figure 4H). As re-expressed ERVs can be detected as pathogens by their host cells^20^, we further analyzed the upregulated proteins of these antiviral pathways and found several effectors of the cytosolic DNA-sensing pathway cGAS-STING (Figure 4K). We validated increased expression of the cGAS-STING pathway components cGAS, STING and RAB2B using western blot, and confirmed increased phosphorylation of the cGAS-STING downstream effector IRF3 in *Setdb1^MuSC-KO^* cells compared to *Control* cells (Figure 4L, S4T). Functional assays using RU521, a specific inhibitor of cGAS, demonstrated a partial rescue of proliferation and cell death in Setdb1*^MuSC-KO^* treated with RU521, confirming the functional consequences of cGAS-STING activation following Setdb1 loss (Figure 4M, 4N). As cGAS-STING is a key mediator of inflammation in many cell types, we suggest it is instrumental to the increased chromatin accessibility (Figure S4J) and increased expression of inflammatory cytokines by *Setdb1^MuSC-KO^* cells (Figure 4B, S4E).

Taken together, multi-omics profiling indicates that *Setdb1* loss-of-function in MuSCs leads to chromatin opening at ERV loci. De-repression of ERVs activates cGAS-STING pathway in *Setdb1^MuSC-KO^* MuSCs leading to cell death and increased cytokine expression.

### SETDB1 in MuSCs regulates regenerative inflammation

To evaluate if aberrant cytokine expression by *Setdb1^MuSC-KO^* MuSCs impact the whole muscle environment, we performed qRT-qPCR for well-known genes involved in cytokine storm on tissue samples.^21^ We found *Setdb1^MuSC-KO^* muscles have higher expression of *Il10*, *Il6*, *Ifng*, *Ccl5*, *Ccl10*, *Ccl2* compared to *Control* muscles at 4 and 7 d.p.i. (Figure 5A, 5B).

**Figure 5.**
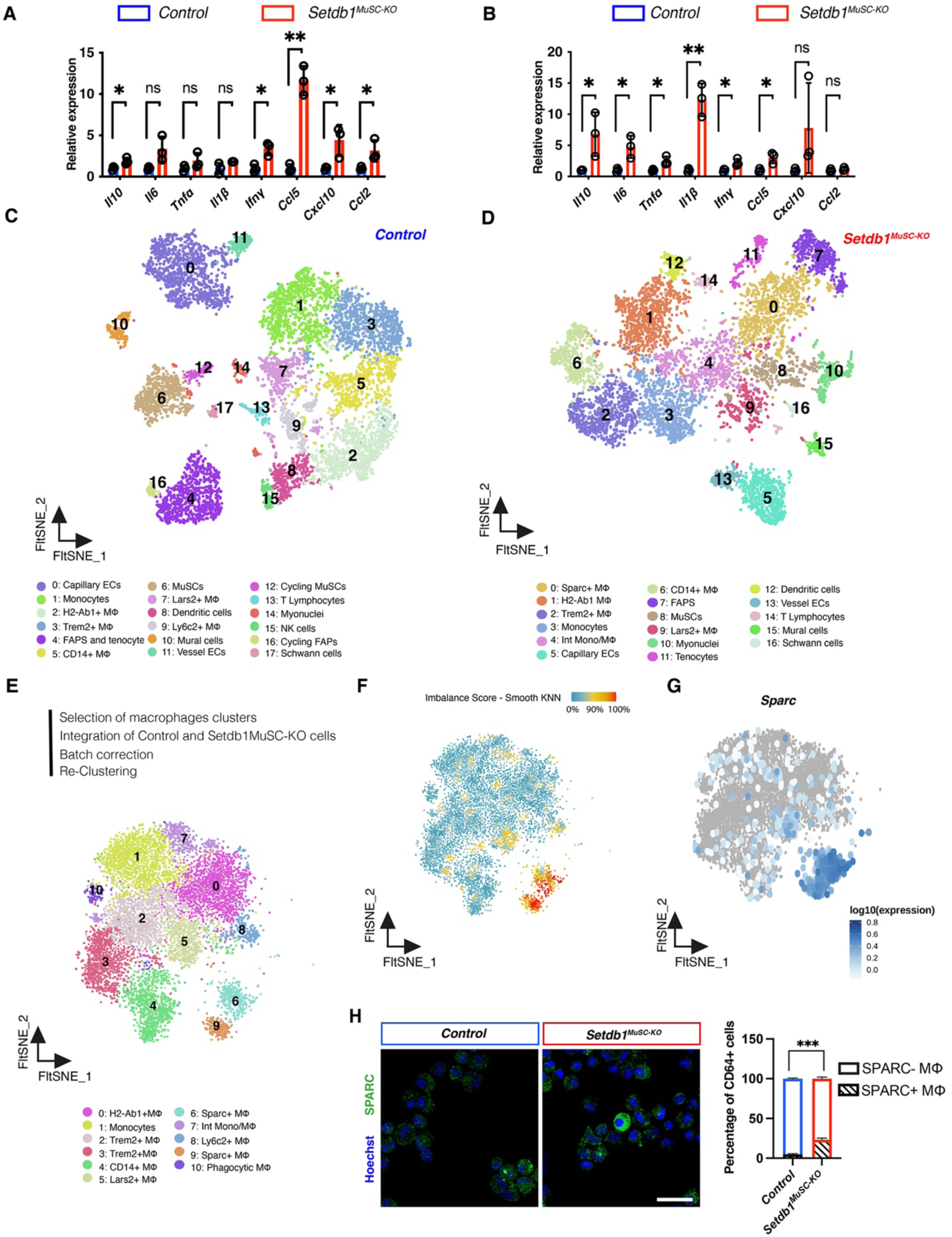
SETDB1 regulate MuSCs signaling toward inflammatory cells. A. RT-qPCR of cytokines and interleukines involved in cytokine storm in *Control* and *Setdb1^MuSC-KO^* TA (*Tibialis Anterior*) cryosections at 4 days after muscle injury using cardiotoxin (CTX). B. RT-qPCR of cytokines and interleukines involved in cytokine storm in *Control* and *Setdb1^MuSC-KO^* cells TA cryosections at 7 days after muscle injury (n=3). C. Single-cell transcriptome analysis of whole muscle in regeneration were isolated from CTX injured hind-limb muscles at 4 days post-injury, and subjected to scRNA-Seq using the 10x genomic platform. The different cell population were computed using Harmony + Louvain algorithm and projected on a FltSNE dimensional reduction, which identified 21 different clusters on separated sample analysis for *Control* samples (n=3 mice pooled in 1 sample). D. Single-cell transcriptome analysis of whole muscle in regeneration were isolated from CTX injured hind-limb muscles at 4 days post-injury, and subjected to scRNA-Seq using the 10x genomic platform. The different cell population were computed using Harmony + Louvain algorithm and projected on a FltSNE dimensional reduction, which identified 15 different clusters on separated sample analysis for *Setdb1^MuSC-KO^* samples (n=3 mice pooled in 1 sample). E. Single-cell transcriptome analysis of macrophages population from *Control* and *Setdb1^MuSC-KO^* samples Dataset were integrated by SCTtransform and different cell population were computed using Harmony + Louvain algorithm and projected on a FltSNE dimensional reduction, which identified 10 different macrophages clusters. F. Density mapping of *Control* and *Setdb1^MuSC-KO^* macrophages population whether nearby cells have the same condition of not. Red regions indicate specific population for the clusters 0 and 9 in *Setdb1^MuSC-KO^*. Blue regions indicate equivalent proportion of population for each sample. G. Single-cell transcriptome analysis of macrophages population from *Control* and *Setdb1^MuSC-KO^* samples Dataset showing the expression pattern of Sparc. Cells are color-coded according to the intensity of Sparc shown. H. Immunostaining for SPARC in *Control* and *Setdb1^MuSC-KO^* macrophages at 4 days after injury after cytospin and quantification of the number of SPARC+ macrophages (n=3 mice). Data are represented as mean ± SD; **p < 0.01, ***p < 0.001, ****p < 0.0001, t test. Scale bars, (H) 50 μm

We next aimed to evaluate the impact of *Setdb1-null* MuSCs on the other cell types residing within the regenerating tissue. To this aim, we performed single-cell RNA-sequencing (scRNA-seq) on TA muscle samples at 4 d.p.i.. We sequenced 10,649 single cells from *Setdb1^MuSC-KO^* samples and 8,169 single cells from *Control* samples. Following normalization, quality control and random resampling, we selected 8,169 single cells for each sample with an average of 9,376 genes expressed per cell in the *Control* condition and 6,494 in the *Setdb1^MuSC-KO^* condition. The scRNA-seq data were analyzed using the R package Seurat.^22^ We used unsupervised graph-based clustering^23^ and embedded cells with FFT-accelerated Interpolation-based t-SNE (Flt-SNE). We annotated clusters to cell types based on differentially expressed genes and patterns of well-described canonical markers (Figure S5A, Figure S5B).

We found single cells from *Control* muscles to cluster adequately into 18 cell types, 9 of which were immune cells (Figure 5C). Among the non-immune cells, we found the excepted muscle-resident cells such as MuSCs, different types of endothelial cells (ECs), fibroblasts (containing both FAPs and tenocytes) and Schwann cells. Remarkably, cell clustering was less efficient on single cells from *Setdb1^MuSC-KO^* muscles. Evaluation of discriminating gene expressions by clusters revealed an inflammatory response signature accompanied by reduced expression of cell-specific markers (Table 4). Furthermore, 17 cell types were identified in the *Setdb1^MuSC-KO^* dataset, with immune cell expansion, reduced MuSC abundance, and FAPs and tenocytes appearing separated, compared to *Control* dataset (Figure 5D). Most interestingly, while both samples contained similar macrophage subtypes (*Lars2+*, *Trem2+*, *H2-Ab1*+, *Cd14+*), *Setdb1^MuSC-KO^* clustering separated the monocytes into two clusters (monocytes and intermediate monocyte/macrophage) and revealed an abundant population of *Sparc*+ macrophages (Figure 5D). To visualize sample-specific cellular subsets, we integrated both datasets and performed clustering. We identified 25 cell types within the integrated map (Figure S5C), as the increase in cell quantity revealed sub-clustering within already identified populations (such as endothelial cells, mural cells, macrophages). Visualization of the single cells sample-of-origin highlighted the cluster 8, namely *Sparc*+ immune cells, as specifically composed only of cells from *Setdb1^MuSC-KO^* muscles (Figure S5D). We next validated the increase abundance of macrophages at 4 d.p.i. in *Setdb1^MuSC-KO^* muscles by flow cytometry (Figure S5E). This augmentation was not associated with a shift in macrophage classical polarization (Figure S5F).

To study the cells appearing in *Setdb1^MuSC-KO^* muscles, we selected and integrated the monocytes and macrophages from both datasets and clustered then anew (Figure 5E). We found 11 distinct populations, including the ones identified in both *Control* and *Setdb1^MuSC-KO^* muscles and the *Sparc*+ population which appeared divided into two sub-clusters (Figure 5E). To precisely map the contribution of each sample to the integrated immune cell dataset, we assigned a score to each single cell based on their origin. This scaling validated the *Sparc^+^* macrophages as specific to regenerating *Setdb1^MuSC-KO^* muscles (Figure 5F), As such, visualization of *Sparc* gene expression onto the integrated immune single cell map showed restricted expression by *Sparc+* macrophages (Figure 5G). We next MACS-sorted macrophages from regenerating *Control* and *Setdb1^MuSC-KO^* muscles to check for SPARC protein expression by immunolocalization of cytospun cells. We found SPARC-expressing Macrophages only in *Setdb1^MuSC-KO^* samples (Figure 5H).

Lastly, we applied Slingshot to infer hierarchical trajectories onto the integrated macrophage populations.^24^ We found two pseudotime lineages both arising from the monocyte cluster. These two lineages encompass populations present in both *Control* and *Setdb1^MuSC-KO^* muscles either progressing through *H2-Ab1+*/*Lars2+* populations or *Trem2+*/*Cd14+* populations to give rise to the *Sparc+* clusters (Figure S5G). Further analysis of the genes differentially expressed along the trajectories suggested increased cell invasion (S100a6, Cald1) and pro-tumoral (Cxcl12, Id1) signatures in *Sparc*+ cells (Figure S5H). Altogether, these data indicate that, in the absence of Setdb1, MuSCs will release cytokines within the regenerating muscle tissue. This is accompanied by accumulation of immune cells in *Setdb1^MuSC-KO^* muscles. Among these, appears a novel *Sparc*+ macrophage lineage with pathological gene expression profile.

### Macrophage histiocytosis and myofiber necrosis follow perturbation of the immune environment in *Setdb1^MuSC-KO^* muscles

The identification of *Sparc+* macrophages prompted us to have a closer look at tissue regeneration in *Setdb1^MuSC-KO^* muscles between the formation of new myofibers and their disappearance (Figure 6A). We observed an impressive 6-fold increase in the macrophage marker F4/80 immunoreactive area in *Setdb1^MuSC-KO^* muscles compared to *Control* muscles at 7 d.p.i. (Figure 6B), Histological analysis of 10 d.p.i. muscles showed a more severe impairment of myofiber formation in *Setdb1^MuSC-KO^* muscles, compared to 7 d.p.i. (Figure 6C). Immunostaining for Lamnin highlighted a complete cellular disorganization of regenerating *Setdb1^MuSC-KO^* muscles (Figure 6D) as well as myofiber atrophy (Figure 6F). By evaluating both F4/80 and Laminin at 10 d.p.i. (Figure 6E), we observed that macrophages were almost absent in control regenerated muscles, while they massively invaded myofibers in *Setdb1^MuSC-KO^* muscles (Figure 6G). These data demonstrate that *Setdb1* gene disruption in MuSCs results in macrophage histiocytosis.

**Figure 6.**
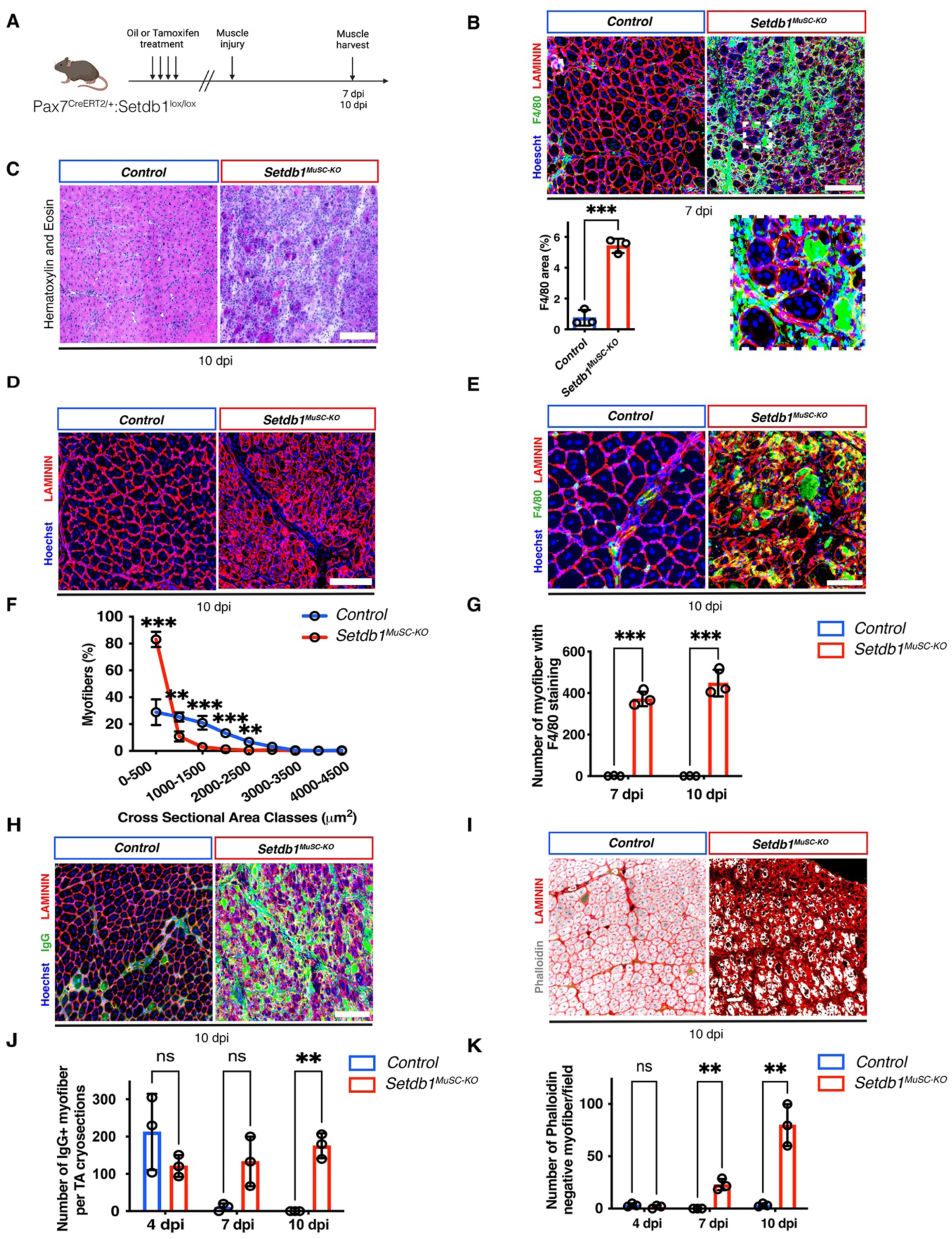
Dysregulation of the inflammatory muscle environment and myofiber necrosis are observed at 7 and 10 dpi in absence of SETDB1 in MuSCs. A. Experimental set up; *Pax7^CRE-ERT2^; Setdb1^fl/fl^* mice intraperitoneally injected with either Oil or Tamoxifen. Cardiotoxin-induced injury (CTX) was performed in the TA (*Tibialis Anterior*) muscles and analyzed 10 post-injury. B. F4/80 and LAMININ immunostaining in *Control* and *Setdb1^MuSC-KO^* muscles at 7 days after injury and quantification of the number of F4/80+ myofibers per field (n=3 mice). C. Hematoxylin and eosin stain of TA muscle cross sections at 10 days post injury (CTX) (n=3 mice). D. LAMININ immunostaining in *Control* and *Setdb1^MuSC-KO^* muscles at 10 days after injury (n=3 mice). E. F4/80 and LAMININ immunostaining in *Control* and *Setdb1^MuSC-KO^* muscles at 10 days after injury and quantification of the number of F4/80+ myofibers per field (n=3 mice). F. Distribution of cross-sectional area of CTX-injured TA muscles in *Control* and *Setdb1^MuSC-KO^* muscles (n=3 mice). G. Quantification of the number of myofibers with F4/80 immunostaining in the *Control* and *Setdb1^MuSC-KO^* muscles at 7 and 10 days post injury (n=3 mice). H. IgG immunostaining in *Control* and *Setdb1^MuSC-KO^* muscles at 10 days after injury (n=3 mice). I. Phalloidin immunostaining in *Control* and *Setdb1^MuSC-KO^* muscles at 10 days after injury (n=3 mice). J. Quantification of the number of IgG+ myofiber per cryosections in the *Control* and *Setdb1^MuSC-KO^* muscles at 4, 7 and 10 days post injury (n=3 mice). K. Quantification of the number of Phalloidin+ myofiber per cryosections in the *Control* and *Setdb1^MuSC-KO Δ^* muscles at 4, 7 and 10 days post injury (n=3 mice). Data are represented as mean ± SD; **p < 0.01, ***p < 0.001, t test. Scale bars, (B) 150 μm (C) 50 μm (D) 100 μm (E) 25 μm (H) 125 μm (I) 50 μm

Macrophage puncture into myofibers happens following acute damage to clean out the tissue from degenerating cells, but also in some inflammatory myopathies in which the diffuse infiltration of macrophages causes myofiber damage.^25^ We thus performed IgG immunostaining at 7 d.p.i. and 10 d.p.i. to evaluate myofiber integrity (Figure 6H). We also stained 4 d.p.i. muscles as an internal control of the experiment, since necrotic myofibers are present following CTX injury in *Control* regenerating muscles. Almost no IgG+ myofibers were detected in control regenerating muscles at 7 d.p.i. and at 10 d.p.i., yet *Setdb1^MuSC-KO^* muscles exhibited high numbers of permeable myofibers at these time points (Figure 6J). As myofibers undergoing necrosis break their sarcomeres down, we stained sections with phalloidin to reveal filamentous Actin (F-Actin) (Figure 6I). While the entirety of myofibers in control muscles showed homogeneous F-Actin expression at all time points, we found F-Actin-negative myofibers in *Setdb1^MuSC-KO^* muscles at 7 d.p.i. and an increased amount of these necrotic myofibers at 10 d.p.i. (Figure 6K).

Taken together, these observations demonstrate that inflammation mediated by the lack of SETDB1 in MuSCs results in inflammation in the myofiber environment but also in penetrance of macrophages into newly formed myofibers and their concomitant necrosis. This process correlates with the complete absence of regenerated myofibers within *Setdb1^MuSC-KO^* muscles at 28 d.p.i., while new myofibers (albeit poorly formed) are present at earlier timepoints following injury.

### SETDB1 is not essential for muscle resident mesenchymal progenitors

We next investigated if targeted *Setdb1* gene disruption in another muscle-resident progenitor cell type, the FAPs, has a similar impact on regeneration as in MuSCs. To this aim, we crossed the *Setdb1* floxed mice with mice with CreERT2 inserted into the start codon of the *Pdgfra* gene.^26^ We used the tamoxifen-inducible recombination protocol on *Pdgfrα*^CreERT2/+:^*Setdb1*^lox/lox^ mice to obtain control and *Setdb1^FAP-KO^* ^mice^ (Figure 7A).

**Figure 7.**
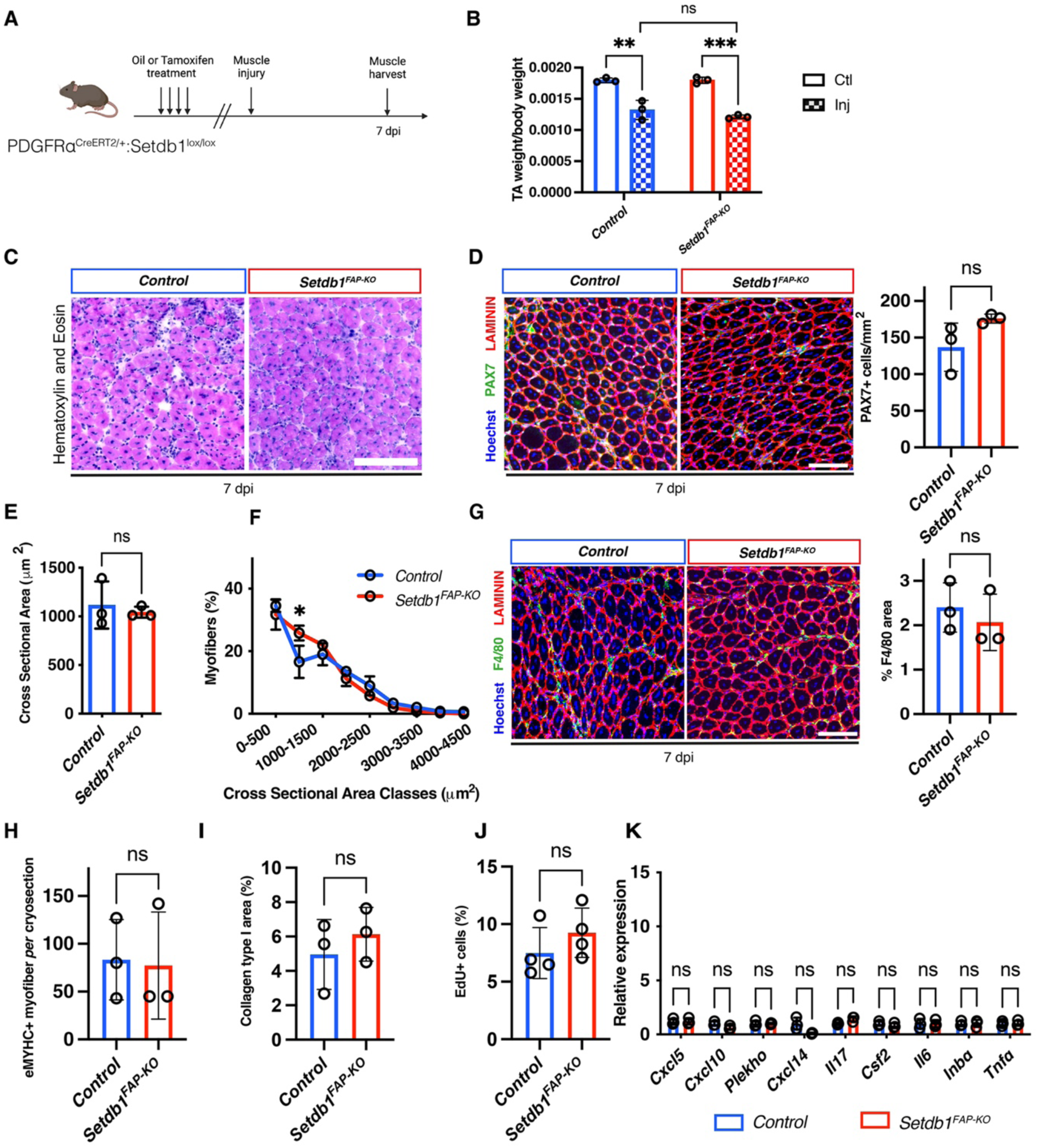
SETDB1 is not essential in muscle resident mesenchymal progenitors. A. Experimental scheme; *PDGFRα^CRE-ERT2^; Setdb1^fl/fl^* mice intraperitoneally injected with either Oil or Tamoxifen and cardiotoxin (CTX) is performed on TA (*Tibialis Anterior*) muscle after 28 days post injury. B. Normalized TA weight on mice body weight represented in *Control* and *Setdb1^FAP-KO^* conditions in uninjured and injured TA (n=3 mice). C. Hematoxylin and eosin stain of TA muscle cross sections at 7 days post injury (CTX) (n=3 mice). D. Immunostaining for LAMININ and PAX7 on TA cryosections 7 days after injury and quantification of the number of PAX7+ cells on TA cryosections (n=3 mice). E. Cross sectional area of CTX-injured TA muscles of *Control* and *Setdb1^FAP-KO^* muscles after 7 days post injury (n=3 mice). F. Distribution of cross-sectional area of CTX-injured TA muscles of *Control* and *Setdb1^FAP-KO^* condition after 7 days post injury (n=3 mice). G. F4/80 and LAMININ immunostaining in *Control* and *Setdb1^FAP-KO^* muscles at 7 days after injury and quantification of the number of F4/80+ myofibers per field (n=3 mice). H. eMYHC+ myofiber per cryosection in *Control* and *Setdb1^FAP-KO^* muscles at 7 days after injury (n=3 mice). I. COLLAGEN I area quantification on TA cryosections 28 days after injury in *Control* and *Setdb1^FAP-KO^* conditions (n=3 mice). J. Quantification of EdU+ cells in *Control* and *Setdb1^FAP-KO^* cells *in vitro* (n=3). K. RT-qPCR on cytokines and interleukins in *Control* and *Setdb1^FAP-KO^* cells *in vitro* (n=3). Data are represented as mean ± SEM; ns not significant, **p < 0.01, t test. Scale bars, (C) 120 μm (D) 80 μm (G) 80 μm.

We then performed CTX injury and did not observe any difference between genotypes in terms of muscle weights (Figure 7B) and muscle histology on cross-sections (Figure 7C) at 7 d.p.i,. Likewise, MuSC quantification at 7 d.p.i. did not reveal any impact of FAP-specific *Setdb1* gene disruption (Figure 7D).

Furthermore, both the number and size of the regenerating myofibers were similar between genotypes (Figure 7E-F). Strikingly, targeted *Setdb1* gene disruption in FAPs neither impacted connective tissue remodeling nor promoted macrophage accumulation in the regenerating muscle (Figure 7H-I). These results demonstrate that *Setdb1* expression is dispensable in FAPs during muscle regeneration.

Finally, we purified FAPs from uninjured muscles of *Pdgfrα*^CreERT2/+:^*Setdb1*^lox/lox^ mice that were treated with Tamoxifen or corn oil *in vivo*. We confirmed efficient *Setdb1* gene deletion *in vitro* (Figure S6A, Figure S6B). We did not observe any difference of the percentage of EdU+ cells between *Control* and *Setdb1^FAP-KO^* cells after 4 days *in vitro* (Figure 7J, Figure S6C). Interestingly, we probed for the expression of the cytokines that were over-expressed by *Setdb1*-null MuSCs and did not find any differences in these gene expression levels between *Control* and *Setdb1^FAP-KO^* FAPs (Figure 7K). Lastly, *Setdb1^FAP-KO^* cells did not show activation of the cGAS-STING pathway (Figures S6D, Figure S6E).

Taken together these data show that, in contrary to MuSCs, SETDB1 has no major role in FAPs and its absence does not result in dysregulation of inflammation *in vivo*.

## DISCUSSION

In this study, we found the KMT SETDB1 is required for MuSCs function. Mechanistically, SETDB1 represses a specific family of transposable elements in MuSCs, the ERVs. We observed that ERVs de-repression in MuSCs leads to MuSCs proliferation arrest and cell death, the activation of cGAS-STING pathway, and entails chemokine release. Moreover, loss of *Setdb1* in MuSCs perturbs the reparative inflammatory response to injury, resulting in macrophage histiocytosis. These dysregulations impair the muscle ability to regenerate and lead to the near complete abolition of the repair process.

We demonstrate SETDB1 maintains MuSC epigenomic landscape integrity, mainly by ensuring chromatin compaction, which we found is necessary for their amplification following exit from quiescence. Without *Setdb1*, MuSCs gradually disappear from the regenerating muscle tissue. Interestingly, *in vitro* studies confirmed that SETDB1 is necessary for MuSC expansion and survival. However, the phenotype of *Setdb1-null* MuSCs appeared more drastic in cell culture than *in vivo*, as these cells were virtually impossible to culture. We speculate the *in vivo* microenvironment is permissive for MuSC survival. As such, during regeneration, MuSCs maintain contacts with the basal lamina, a major component of their niche, and interact with other cell types known to enhance MuSC proliferation/survival such as pro-inflammatory macrophages and activated FAPs. It is also possible cell death signals are concentrated in the *in vitro* environment since only one pure population of myogenic cells is present in the cultures.

Transposable Elements (TEs) are genetic elements with the ability to replicate and transpose within the genome of germline and of specific somatic cells.^27^ Close to 40% of the mouse genome is made up of transposable elements and approximately 10% represent endogenous retrovirus (ERV) sequences. These ERV sequences have resulted from both ancient and modern infections by exogenous retroviruses, which have successfully colonized the germline of their host. As observed in our work, muscle cells express ERVs at the basal level. However, ERVs have been studied and found upregulated and disastrous in context of cancer, autoimmune diseases and neurodegenerative pathologies. Their functions, and especially the consequences of their re-expression, have been poorly studied in the context of adult tissue regeneration. Only one role for ERVs had been established in skeletal muscle, as they may be involved in statin-induced myopathies.^28,29^ In contrast, our results indicate that when MuSCs exit from quiescence, ERVs de-repression has more severe consequences, triggering disaggregation of regenerating muscle fibers. We suggest that active repression of ERVs in muscle cells is a crucial process to allow for normal muscle physiology.

The cGAS-STING cytosolic DNA-sensing pathway and its link with inflammation has been extensively studied these past years.^30–32^ Many works highlighted the activation of the pathway in an anti-viral response context. In addition, cGAS-STING is involved in anti-proliferative process and cell death, notably in mouse embryonic fibroblasts (MEFs) and HeLa cells.^33–35^ Importantly, the link between SETDB1 and the cGAS-STING pathway has been described in two recent studies performed in cancer cells.^18,19^ Here, we show that the activation of the cGAS-STING pathway in *Setdb1-null* MuSCs results in hypercytokinemia and is correlated with an aberrant inflammatory response within *Setdb1^MuSC-KO^* regenerating tissue. It will be an area of future investigation to precisely evaluate if the increased cytokine expression by *Setdb1-null* MuSCs is causal to the observed macrophage histiocytosis. As such, it may be interesting to evaluate the impact of cGAS-STING inhibitors *in vivo* during muscle regeneration.

Further to that point, our results suggest that proliferating *Setdb1-null* MuSCs, with active cGAS-STING pathway, will perturb the orchestration of immune cells infiltrating the wounded tissue. Strikingly, we describe the appearance of a new population of macrophages only in regenerating *Setdb1^MuSC-KO^* muscles. These *Sparc+* macrophages do not present the same features as healthy macrophages required for efficient muscle regeneration.^37^ The inflammatory cytokines released by *Setdb1-null* MuSCs are known to influence monocyte polarization into either pro- or anti-inflammatory macrophages, notably in cancer.^39,41^ Similarly, the cGAS-STING pathway has been shown to be related to the acquisition of tumor-associated-macrophage (TAMs) phenotype.^42,44^ TAMs are well described as a dysregulated immune cell population involved in tumor growth and metastasis and demonstrating an aggressive behavior *in vivo*.^43^ Strikingly, the *Sparc+* macrophages have TAM characteristics such as matrix formation (expression of collagens, fibronectins) and epithelial-mesenchymal-transition. Future studies may test whether these *Sparc+* cells result from failed regeneration of myofibers or are directly causing the necrosis and disaggregation of *Setdb1-null* myofibers.

Finally, we found that targeted deletion of *Setdb1*-targeted deletion in FAPs neither affect their cell proliferation capacity, nor the muscle regeneration process following acute injury. This discrepancy with MuSCs can be explained by a divergence in the actors dedicated to histone methylation by these different muscle-resident progenitor types. Interestingly, targeted disruption of another H3K9 methyltransferase, G9a (KMT1C), in MuSCs has no effect on muscle regeneration.^45^ In contrast, G9a has recently been shown to be required for the maintenance of FAP identity *in vivo*^46^. Taken together, these observations clearly indicate that different stem/progenitor cell types within a given tissue have unequal requirements for specific H3K9 KMTs. One can also expand this concept to propose that different stem/progenitor cell may have divergent needs for TE silencing. Indeed, in mESCs, H3K9 KMTs favors different TE classes, like SUV39H1 (KMT1A), which represses class II ERVs, such as the intracisternal A-particles (IAPs), but also the long interspersed nuclear elements (LINEs) such as LINE1 in mESCs^47^ while SETDB1 is known to be recruited for the repression of the class I and II of Long terminal repeat (LTR) and IAPs in mESCs.^17^ Therefore, our results advocate that SETDB1 may be an important regulator of adult stem cell function in general, but other H3K9 KMTs should be considered depending on tissues or cell types.

In conclusion, we identified the H3K9 KMT SETDB1 to be required for ERVs silencing in adult MuSCs, but not FAPs. Through preventing unscheduled ERV expression during MuSCs following activation from quiescence, SETDB1 allows for their amplification and for proper inflammation process, preventing histiocytosis and myofiber necrosis. These findings represent a significant advance in our understanding of MuSCs biology and muscle regeneration. Future work will investigate the role of SETDB1 in context of pathologies such as inflammatory myopathies and of sarcopenia due to aging.

## ACKNOWLEDGEMENTS

The research conducted in the Le Grand lab was supported by funding from the Inserm, the CNRS, the Agence National pour la Recherche (ANR-19-CE13-0016, ANR-17-CE12-0010-02), the Association Française contre les Myopathies/AFM-Téléthon (MyoNeurAlp2 project) and the European Joint Program for Rare Diseases (EJPRD JTC2019, Myocity project). C.B. was supported by a post-doctoral fellowship from the Fondation pour la Recherche Médicale (FRM). P.G. was supported by a 4rth year of PhD from the FRM. The authors acknowledge the PLATIM and CIQLE microscopy facilities of Université Claude Bernard Lyon 1. We thank C. Degletagne for support with single cell RNA-seq. We thank B. Chazaud and G. Juban for their help on the macrophages work.

## AUTHOR CONTRIBUTIONS

F.L.G and S.A conceived the study, designed the project, and obtained grant funding. F.L.G supervised the project and managed the laboratory. P.G performed most of the experiments. W.J and S.F performed bio-informatic analyses. C.B performed cell biology experiments and helped P.G with in vivo experiments. L.G performed preliminary experiments. C.P setup the mice colony and performed preliminary experiments. F.A and J.R performed cell culture experiments. W.K and T.C performed proteomics. P.G, T.C, S.A and F.L.G designed or interpreted experiments. P.G and F.L.G wrote the manuscript. All authors reviewed and edited the manuscript.

## SUPPLEMENTAL FIGURES

**Supplemental Figure S1, related to Figure 1.**
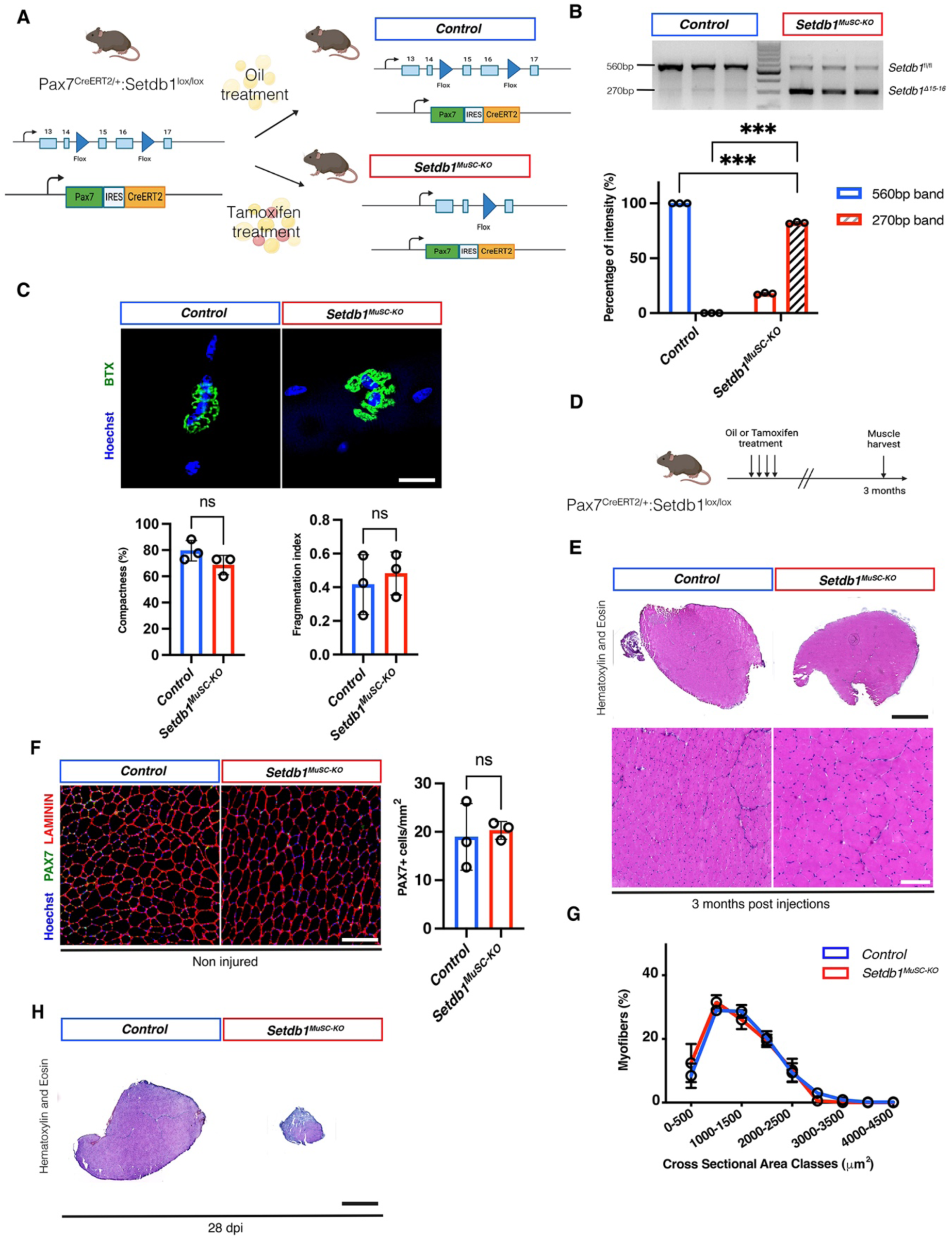
The loss of *Setdb1* does not affect muscle homeostasis. A. Conditional knock-out of *Setdb1* in adult MuSCs using Tamoxifen-inducible CRE under the MuSC specific PAX7 promoter. B. Genotyping of *Control* and *Setdb1^MuSC-KO^* cells *in vitro* and quantification of the band intensity (n=3 mice). C. Immunostaining for Bungarotoxin on single myofiber fixed in PFA at time 0h after digestion in *Control* and *Setdb1^MuSC-KO^* and quantification of the compactness and fragmentation index in both conditions (n=3). D. Experimental scheme; *Pax7^CRE-ERT2^; Setdb1^fl/fl^* mice intraperitoneally injected with either Oil or Tamoxifen and are harvested 3 months after the injections. E. Hematoxylin and eosin stain of TA (*Tibialis Anterior*) muscle cross sections at 3 months after oil and tamoxifen injections (n=3 mice). F. Quantification of PAX7+ cells in *Setdb1^+/+^* and *Control* and *Setdb1^MuSC-KO^* muscles *in vivo* at 3 months after oil and tamoxifen injections (n=3 mice). G. Distribution of cross sectional area of TA muscles of *Control* and *Setdb1^MuSC-KO^* TA muscles at 3 months after injections. H. Hematoxylin and eosin stain of TA muscle cross sections at 28 days post injury (CTX) (n=3 mice). Data are represented as mean ± SD; ns not significant, ***p < 0.001, t test. Scale bar, (C) 20μm, (E) 50 μm, (F) 50 μm, (G) 200μm.

**Supplemental Figure S2, related to Figure 2.**
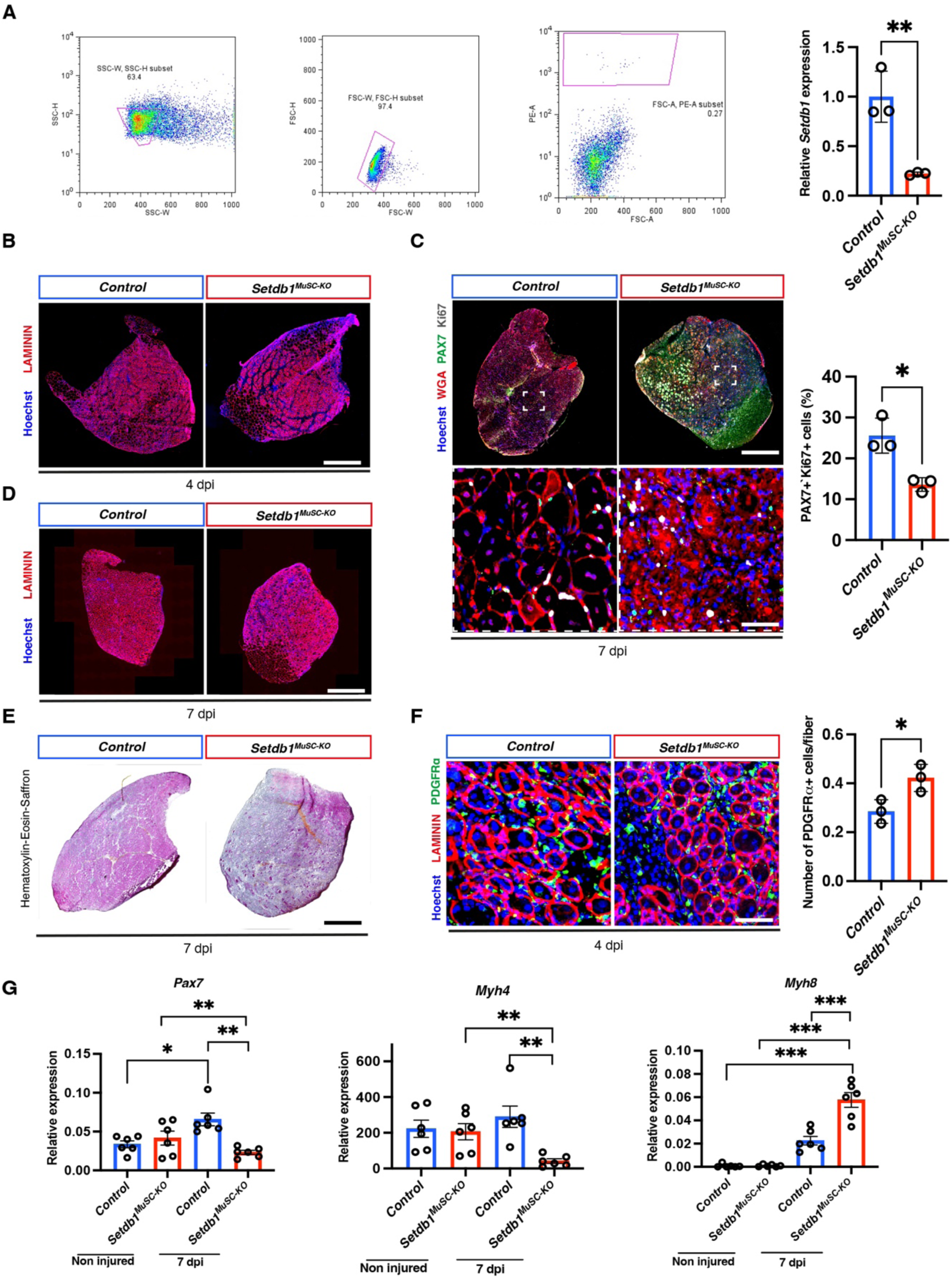
The muscle regeneration process is affected by the loss of *Setdb1* at 4 d.p.i. and 7 d.p.i.. A. Relative expression of *Setdb1* on MuSCs-FACS sorted cells using the Tdt reporter (n=3). Immunostaining against LAMININ in TA (*Tibialis Anterior*) muscle cross sections after cardiotoxin (CTX) injection at 4 days after injury in *Control* and *Setdb1^MuSC-KO^* conditions (n=3 mice). B. Immunostaining against LAMININ on whole cryosections after 4 days post injury in the *Control* and *Setdb1^MuSC-KO^* condition. C. Immunostaining against PDGFRα in TA muscle cross sections after CTX injection at 4 days after injury in *Control* and *Setdb1^MuSC-KO^* muscles and quantification of the number of PDGFRα+ cells (n=3 mice). D. Immunostaining against LAMININ in TA muscle cross sections after CTX injection at 7 days after injury in *Control* and *Setdb1^MuSC-KO^* muscles (n=3). E. Immunostaining against Ki67, PAX7 and LAMININ in TA muscle cross sections after CTX injection and quantification of the PAX7+Ki67+ cells at 7 days after injury in *Control* and *Setdb1^MuSC-KO^* muscles (n=3). F. Hematoxylin-Eosin-Saffron staining on TA cryosections at 7dpi (n=3). G. RT-qPCR of Pax7, Myh4 and Myh8 on TA cryosections at uninjured or 7 days after CTX in *Control* and *Setdb1^MuSC-KO^* muscles (n=6). Data are represented as mean ± SD; ns not significant, *p < 0.05, **p < 0.01, ****p < 0.0001. Scale bars, (B) 200 μm, (C) 200 μm, 25μm (D) 200 μm, (E) 200 μm (F) 50 μm.

**Supplemental Figure S3, related to Figure 3.**
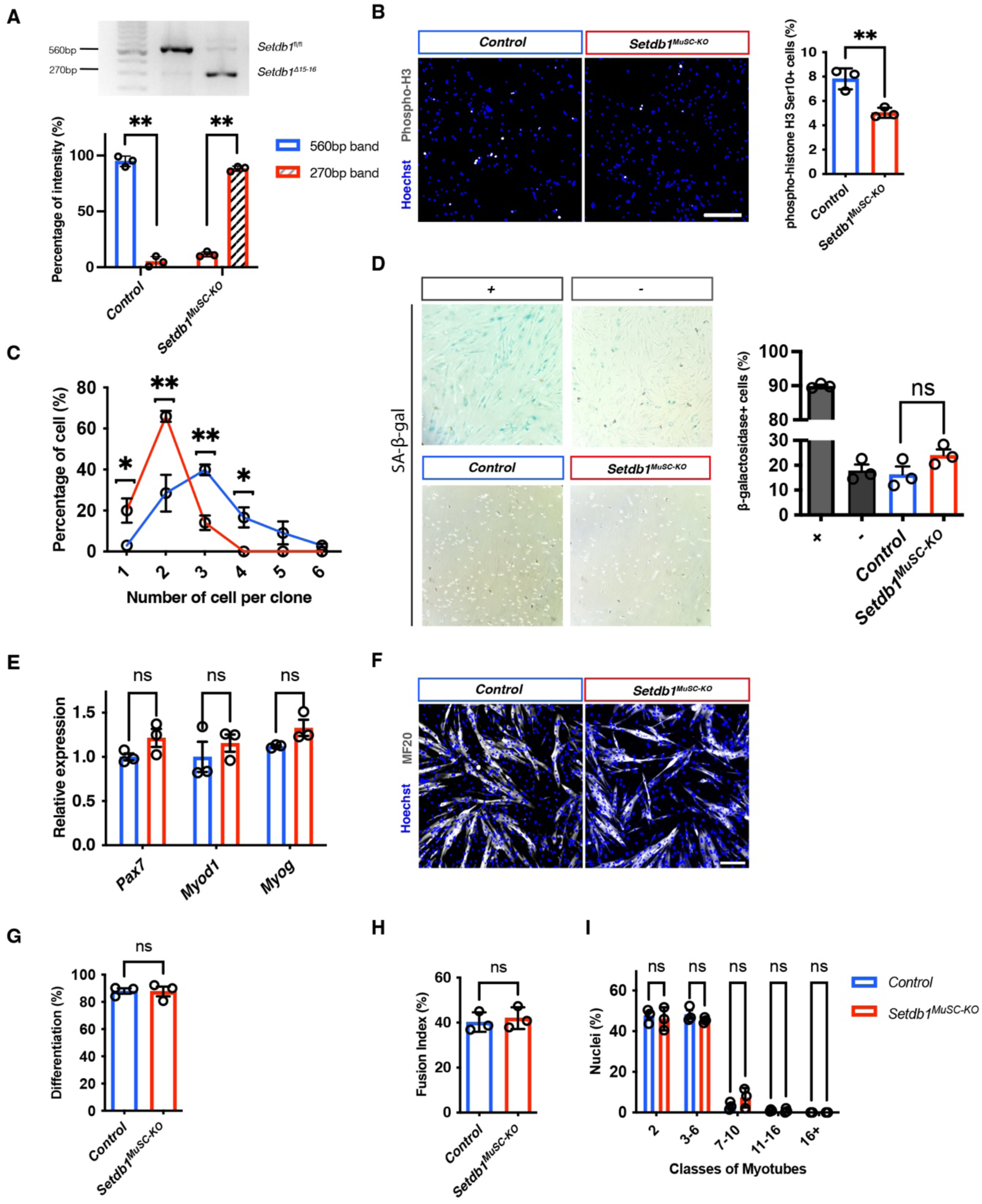
The loss of *Setdb1* promotes cell cycle defect but does not impact muscle differentiation *in vitro*. A. Clonal assay showing the number of cell per clone after 8 days in *Control* and *Setdb1^MuSC-KO^ condition* (n=3). B. Genotyping and quantification showing *Setdb1^fl/fl^* deleted after 5 days of ethanol or 4-hydroxitamoxifen treatment. C. Immunostaining against phospho-Histone 3 *in vitro* after 5 days of ethanol or 4-hydroxitamoxifen treatment and quantification of the phospho-histone H3+ cells (n=3). D. β-galactosidase assay performed *in vitro* after 5 days of ethanol or 4-hydroxitamoxifen treatment showing the quantification of the β-galactosidase positive cells (n=3). E. Relative expression of *Pax7*, *Myod1* and *Myog* in MuSCs *in vitro* after 5 days of ethanol or 4-hydroxitamoxifen treatment (n=3). F. Immunostaining against MF20 in *Control* and *Setdb1^MuSC-KO^* cells *in vitro* after 3 days of differentiation (n=3). G. Quantification of the percentage of differentiated cells in *Control* and *Setdb1^MuSC-KO^* conditions (n=3). H. Quantification of the percentage of fused cells in *Control* and *Setdb1^MuSC-KO^* conditions (n=3). I. Proportion of the number of nuclei in the *Control* and *Setdb1^MuSC-KO^* conditions *in vitro* (n=3). Data are represented as mean ± SD; ns not significant, **p < 0.01, t test. Scale bars, (B) 75 μm, (D) 200um (F) 75 μm.

**Supplemental Figure S4, related to Figure 4.**
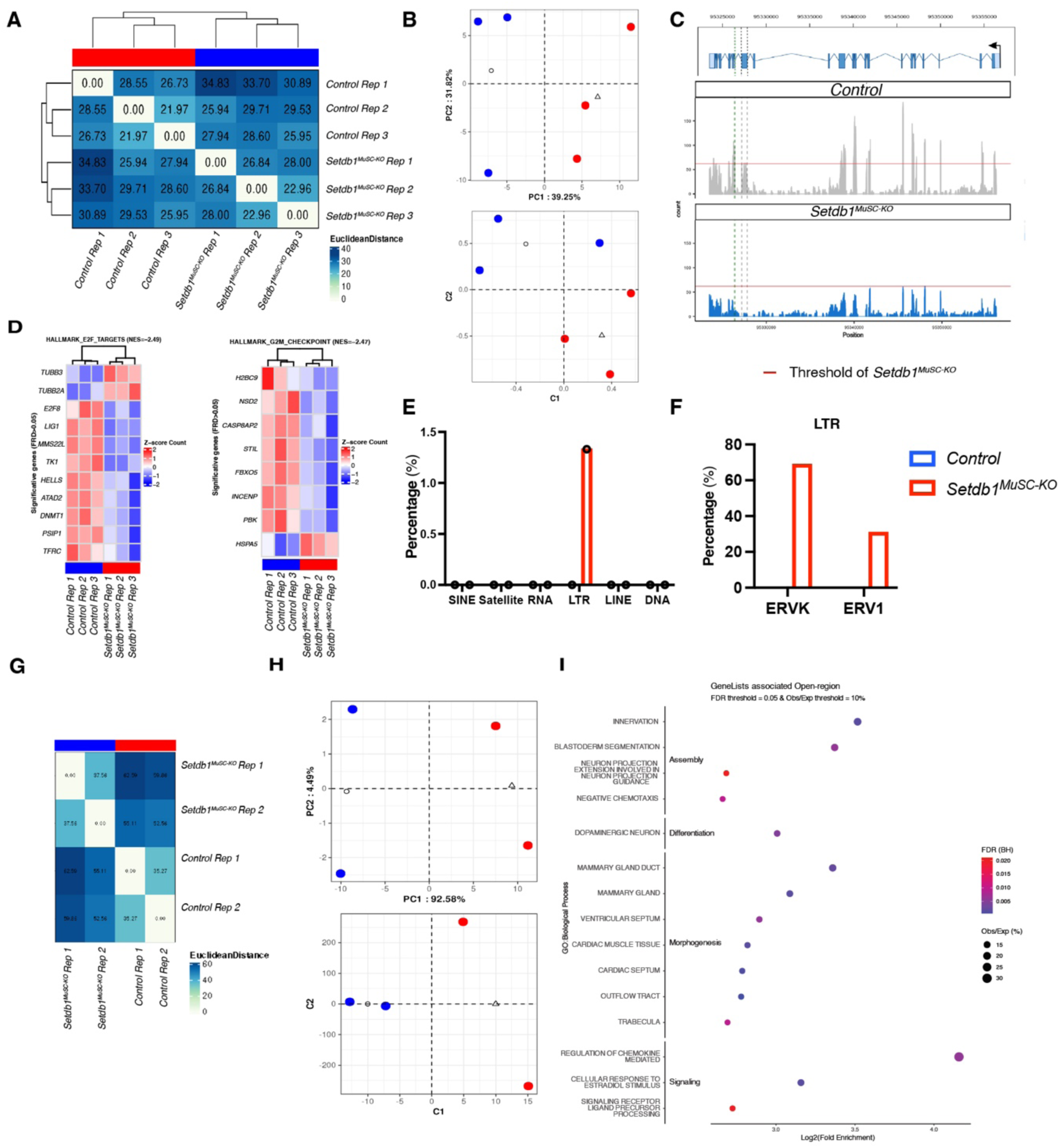
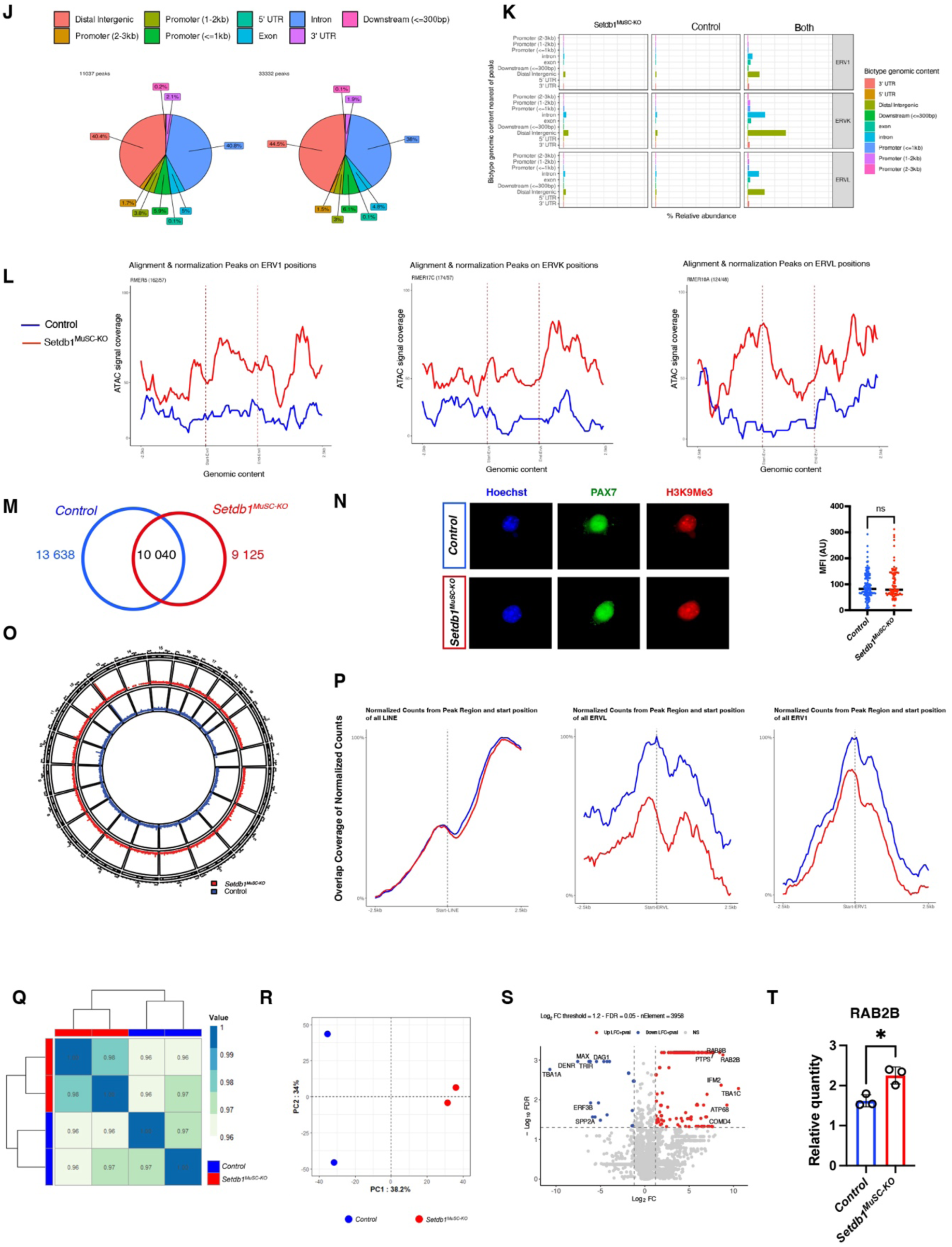
The loss of *Setdb1* promotes ERV de-repression and inflammation. A. Visualizing the overall effect of experimental covariates and batch effects using first (PC1) and second (PC2) Principal Component Analysis (PCA) dimension (in top) and first (C1) and second (C2) Uniform Manifold Approximation and Projection (UMAP) dimensions (in bottom) on the 1000 most variable expression genes. B. Unsupervised clustering of the RNA-seq samples over similarities and dissimilarities by sample from euclidean distance matrix on the 1000 most variable expression genes. C. Tracks of reads mapped on Setdb1 gene (ENSMUSG00000015697) superposed on canonical transcript (ENSMUST00000015841,12).in the *Control* and *Setdb1^MuSC-KO^* cells. D. Heatmap of RNA-seq normalized expression z-scores computed on significative genes (FDR < 0.05) for hallmark E2F target and G2M checkpoint in the *Control* and *Setdb1^MuSC-KO^* cells (n=3 mice). Genes and samples are clustered using euclidean matrix. E. Percentage of the different transposable element families found in the *Control* and *Setdb1^MuSC-KO^* cells. F. Percentage of the different Long Terminal Repeat found in the *Control* and *Setdb1^MuSC-KO^* cells. G. Visualizing the overall effect of experimental covariates and batch effects using first (PC1) and second (PC2) Principal Component Analysis (PCA) dimension (in top) and first (C1) and second (C2) Uniform Manifold Approximation and Projection (UMAP) dimensions (in bottom) on the 1000 most variable expression peaks. H. Unsupervised clustering of the ATAC-seq samples over similarities and dissimilarities by sample from euclidean distance matrix on the 1000 most variable expression genes. I. Functional enrichment analysis directly performed on genomic regions by rGREAT tool on open-region between *Control* and *Setdb1^MuSC-KO^* (n=2). Dot in red indicate a high significant enrichment (adjusted by FDR), dot in blue indicate a low significant enrichment. The size shows the number of open-region associated to gene in process (observation) on the number of open-region associated to all gene in the same process (expected). Results are ordering in high to low level of Log2 Fold Change. Selected enriched terms are presented according to the significativity of results (FDR < 0,05) and ratio observation/expected >= 10%. J. Pie plot representing the different type of genomic regions opened in the *Control* and *Setdb1^MuSC-KO^* cells. K. Relative abundance of the different locations of the ERV found in the *Control* and *Setdb1^MuSC-KO^* cells inside each genomic regions opened. L. Footprinting K-mean of the different ERV families found in the Control and *Setdb1^MuSC-KO^* cells. M. Venn diagram representing the peak number found in the H3K9Me3 chip-seq in the *Control*, *Setdb1^Musc-KO^* and both conditions (n=1). N. PAX7 and H3K9Me3 immunostaining on Control and Setdb1MuSC-KO cells in vitro and quantification of the mean fluorescence intensity of the H3K9Me3 staining (n=3). O. Genomic distribution of H3K9Me3 Chip-seq peaks (log10 (peak score) > 1.2) on mouse genome for *Control* and *Setdb1^MuSC-KO^* cells *in vitro* (n=1). P. Overlap coverage of normalized counts from H3K9Me3 Peak Region and start position of all LINE, ERVL and ERV1 in both conditions. Q. Unsupervised clustering of the mass spectrometry samples over similarities and dissimilarities by sample from euclidean distance matrix on the 1000 most variable expression protein. R. Visualizing the overall effect of experimental covariates and batch effects using first (PC1) and second (PC2) Principal Component Analysis (PCA) on the 1000 most variable expression proteins (n=2). S. Volcano plot over *Setdb1^MuSC-KO^* muscles mass spectrometry data (n=2). Dot in colors indicate significant dysregulated proteins (FDR<0.05): in red, up-dysregulated (Log2 Fold-Change (LFC)>=1), in blue down-dysregulated (LFC<=-1). Dot in grey indicate no significant dysregulated proteins or low value of LFC (abs(LFC)<1). T. Western blot quantification of RAB2B in both conditions (n=3). Data are represented as mean ± SD; ns not significant, *p < 0.05, **p < 0.01, t test.

**Supplemental Figure S5, related to Figure 5.**
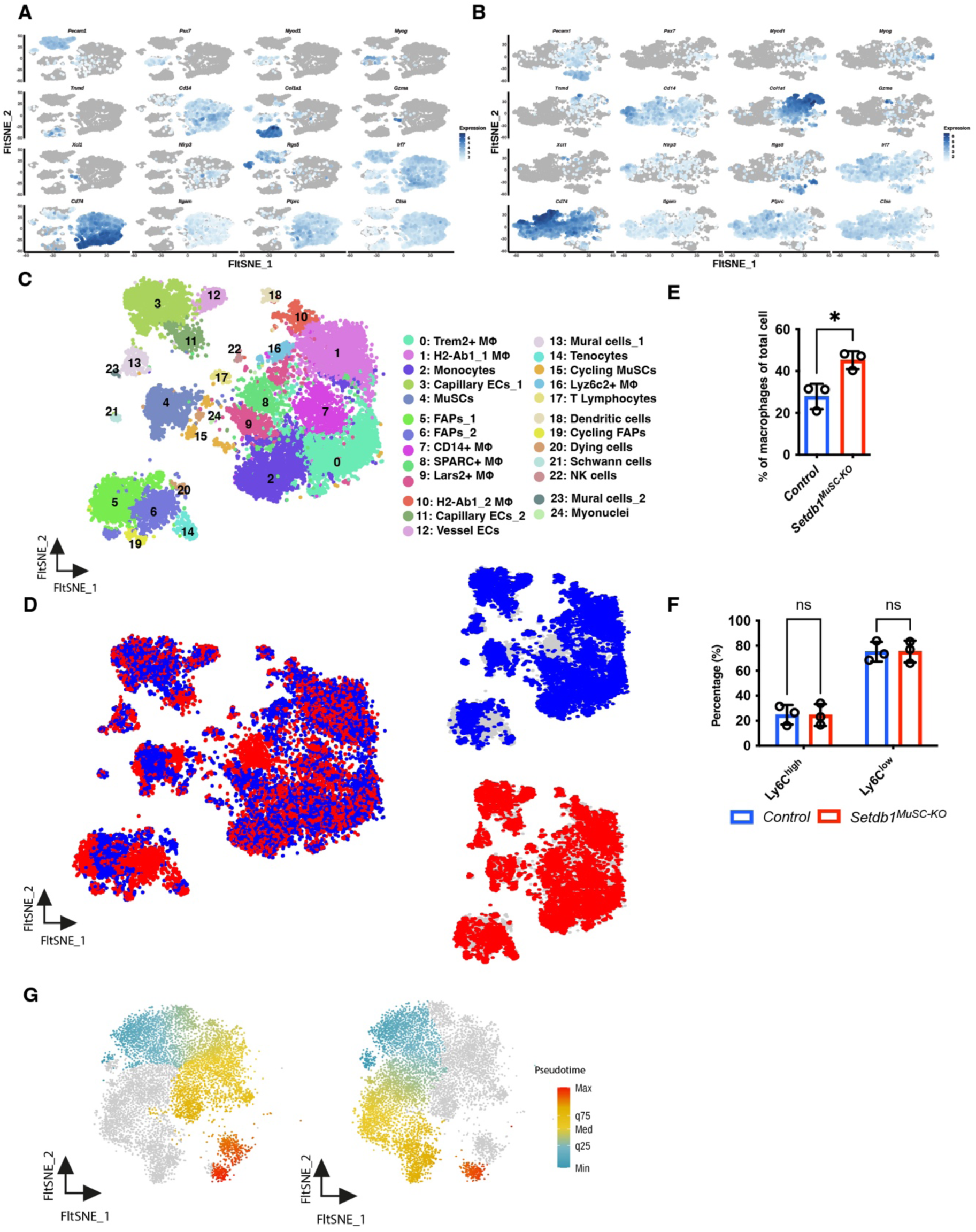
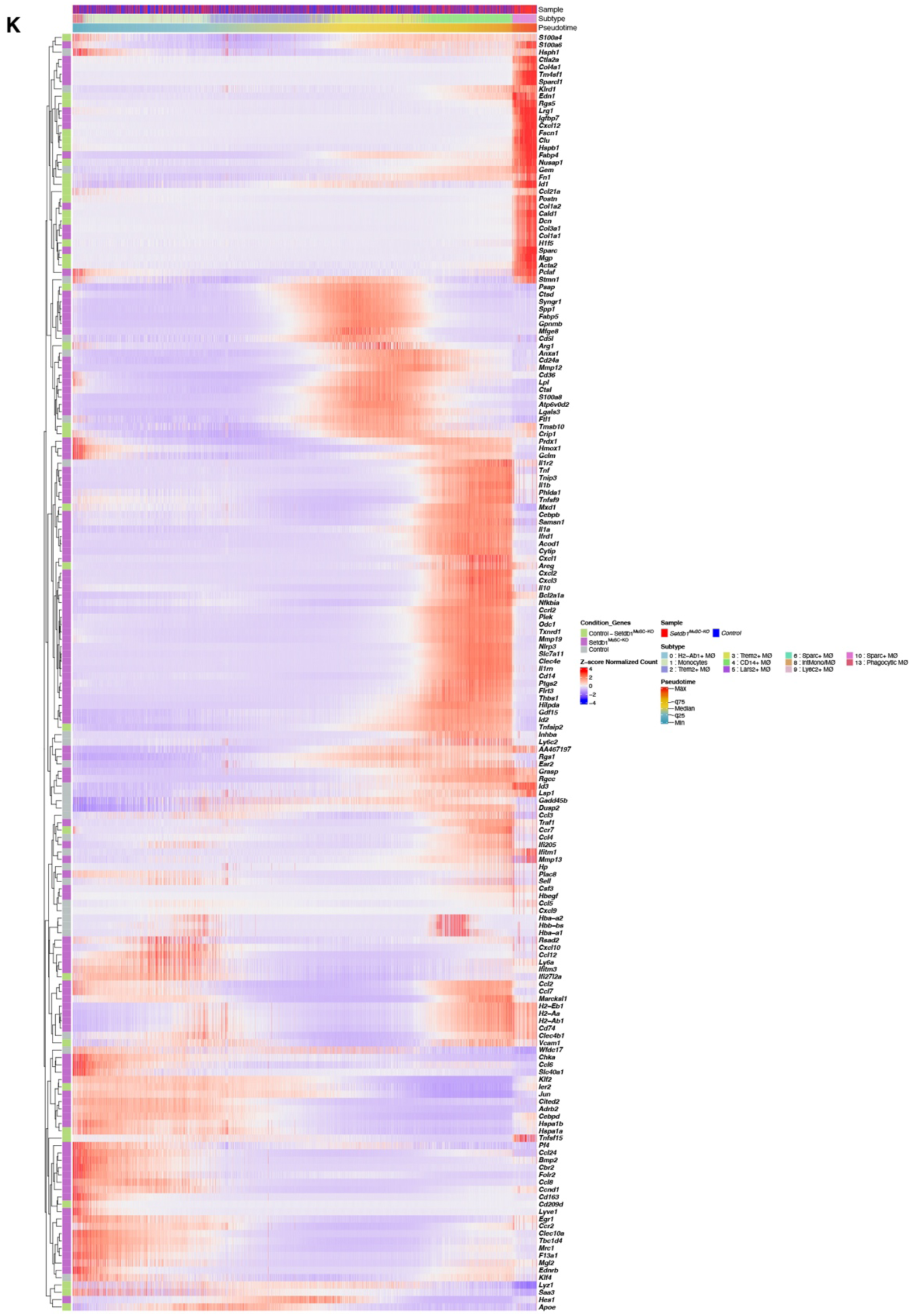
Single cell RNA-seq analysis of 4 d.p.i. regenerating muscles. A. Feature Plot projected on a FltSNE dimensional reduction of the scRNA-seq from the *Control* condition. Each graph shows the expression pattern of a selected canonical marker used to identify the different populations. Cells are color-coded according to the intensity of the marker shown. B. Feature Plot projected on a FltSNE dimensional reduction of the scRNA-seq from the *Setdb1^MuSC-KO^* condition. Each graph shows the expression pattern of a selected canonical marker used to identify the different populations. Cells are color-coded according to the intensity of the marker shown. C. Single-cell transcriptome analysis of whole muscle in regeneration were isolated from CTX injured hind-limb muscles at 4 days post-injury, and subjected to scRNA-Seq using the 10x genomic platform. The different cell population were computed using Harmony + Louvain algorithm and projected on a FltSNE dimensional reduction, which identified 25 different clusters on separated sample analysis for *Control* and *Setdb1^MuSC-KO^* integrated samples (n=3 mice pooled in 1 sample). D. Single-cell transcriptome analysis showing the integrated FltSNE with both *Control* and Setdb1*^MuSC-KO^*. E. Percentage of macrophages found in *Control* and *Setdb1^MuSC-KO^* mice using flow cytometry at 4 days after TA (*Tibialis Anterior*) injury using cardiotoxin (CTX) (n=3 mice). F. Percentage of pro and anti-inflammatory macrophages found in *Control* and *Setdb1^MuSC-KO^* mice using flow cytometry at 4 days after TA injury using CTX (n=3). G. Pseudotime trajectory using the Slingshot algorithm shows the cell trajectory in 4 projected lines across 10 clusters from the *Fol2*+ monocytes to the different populations of macrophages. H. Heatmap of top100 high variables genes differentially expressed along the fourth calculated trajectory of *Setdb1^MuSC-KO^* condition. Data are represented as mean ± SD; ns not significant, *p < 0.05, t test.

**Supplemental Figure S6, related to Figure 7.**
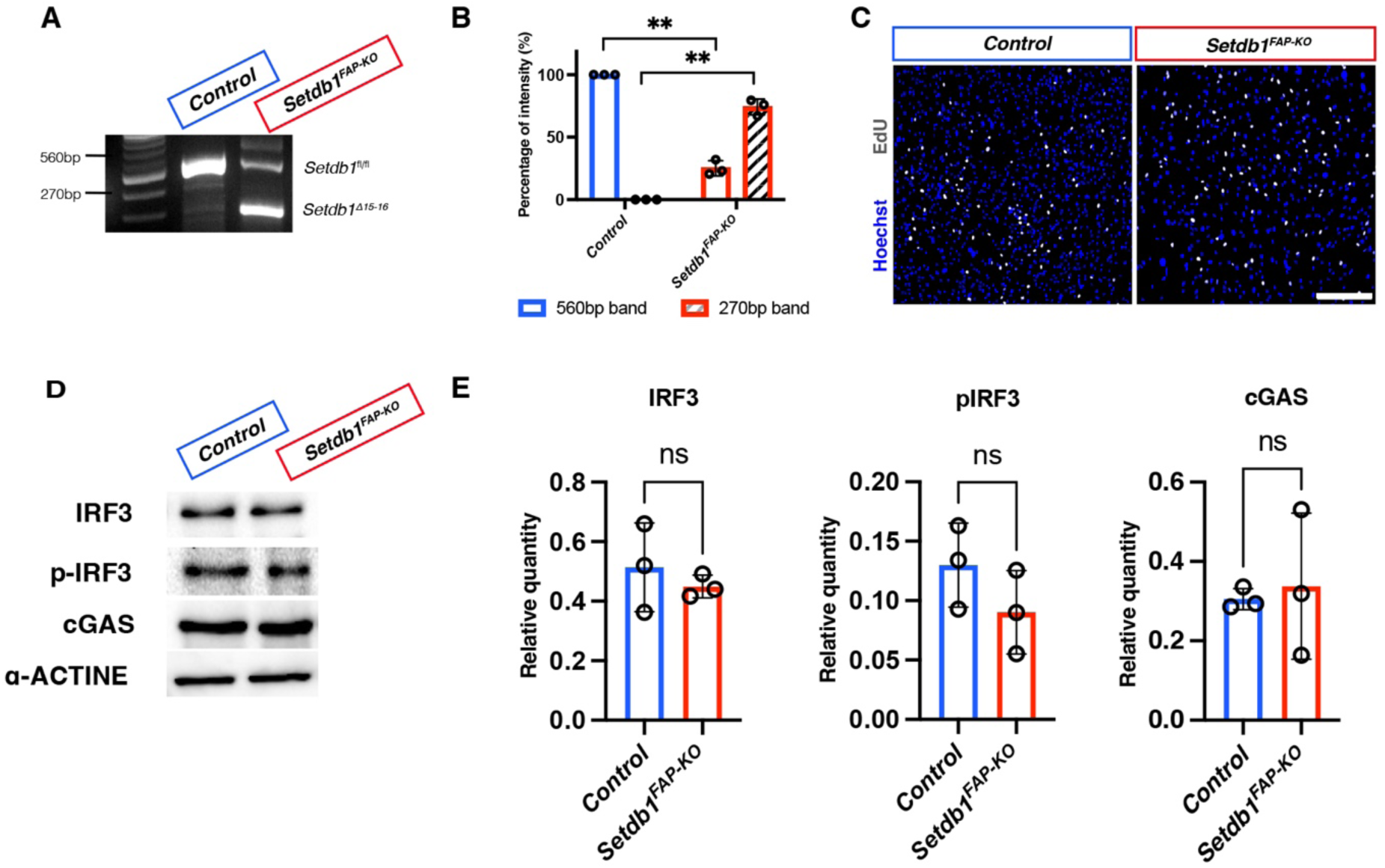
The loss of *Setdb1* in FAPs does not affect muscle regeneration *in vivo*. A. Genotyping of *Control* and *Setdb1^FAP-KO^* fibroblasts *in vitro* and quantification of the band intensity. B. Quantification of the band intensity of *Control* and *Setdb1^FAP-KO^* fibroblasts *in vitro.* C. Immunostaining against EdU in Control and *Setdb1^FAP-KO^* fibroblasts *in vitro* and quantification of EdU+ cells in both conditions. D. Western blot analysis of IRF3 (55 kDa), p-IRF3 (55 kDa) and cGas (60 kDa) in *Control* and *Setdb1^FAP-KO^* fibroblasts *in vitro* and quantification of the relative quantity. E. Western blot quantification of IRF3, p-IRF3 and cGas in *Control* and *Setdb1^FAP-KO^* fibroblasts *in vitro* and quantification of the relative quantity. Data are represented as mean ± SD; ns not significant, *p < 0.05, **p < 0.01, t test. Scale bar, (C) 150 μm.

## METHODS

### KEY RESSOURCES TABLE

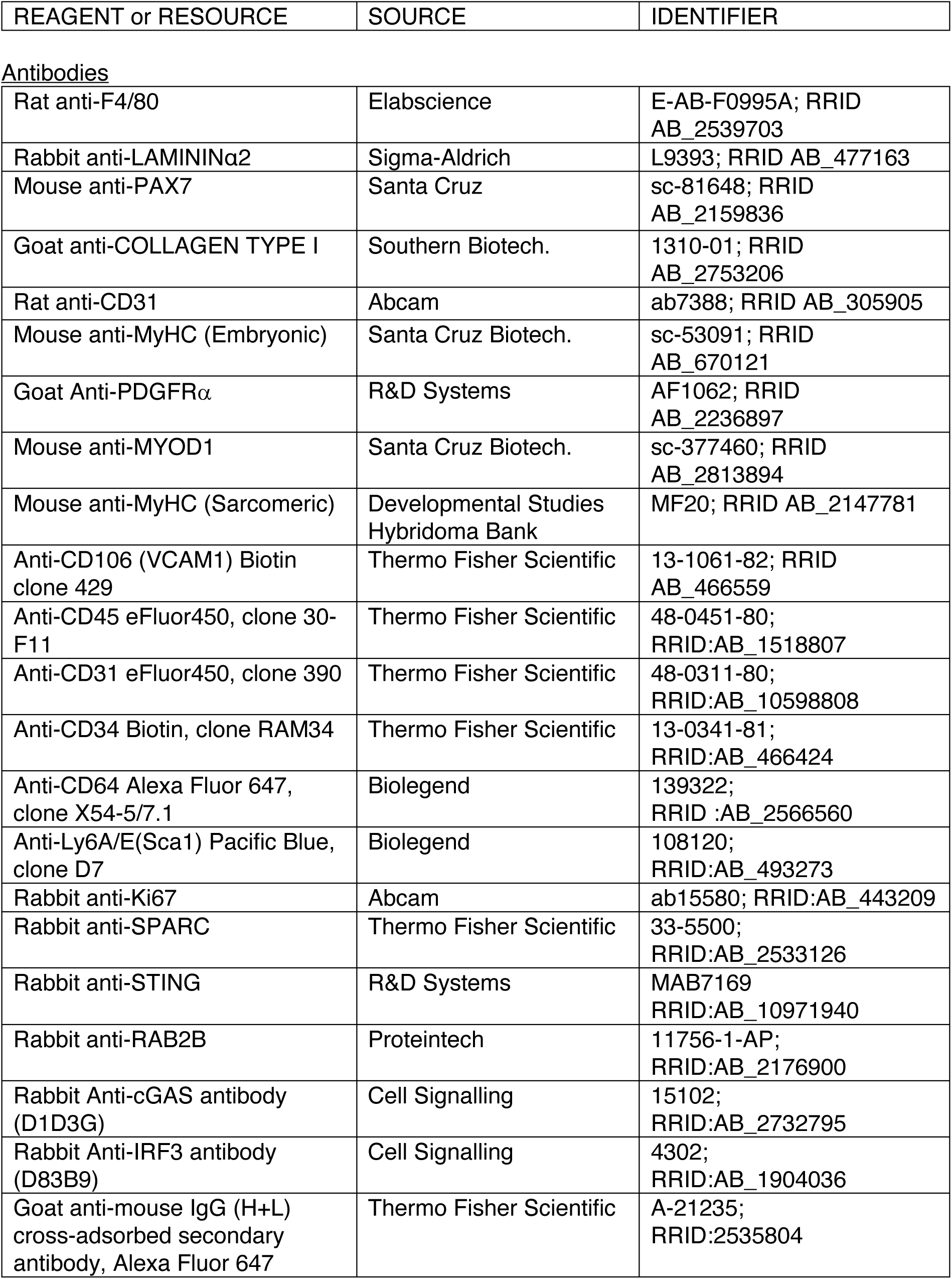
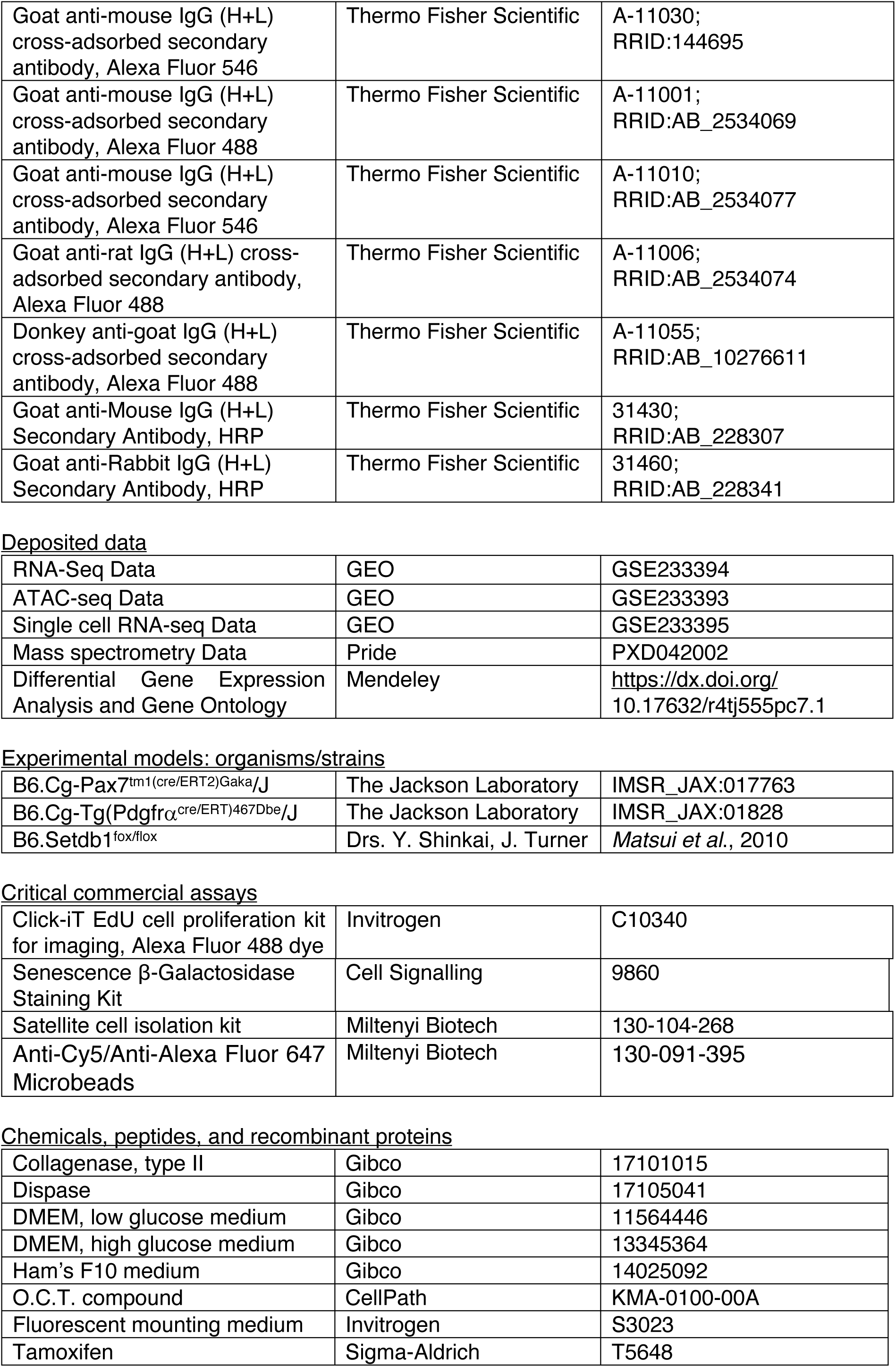
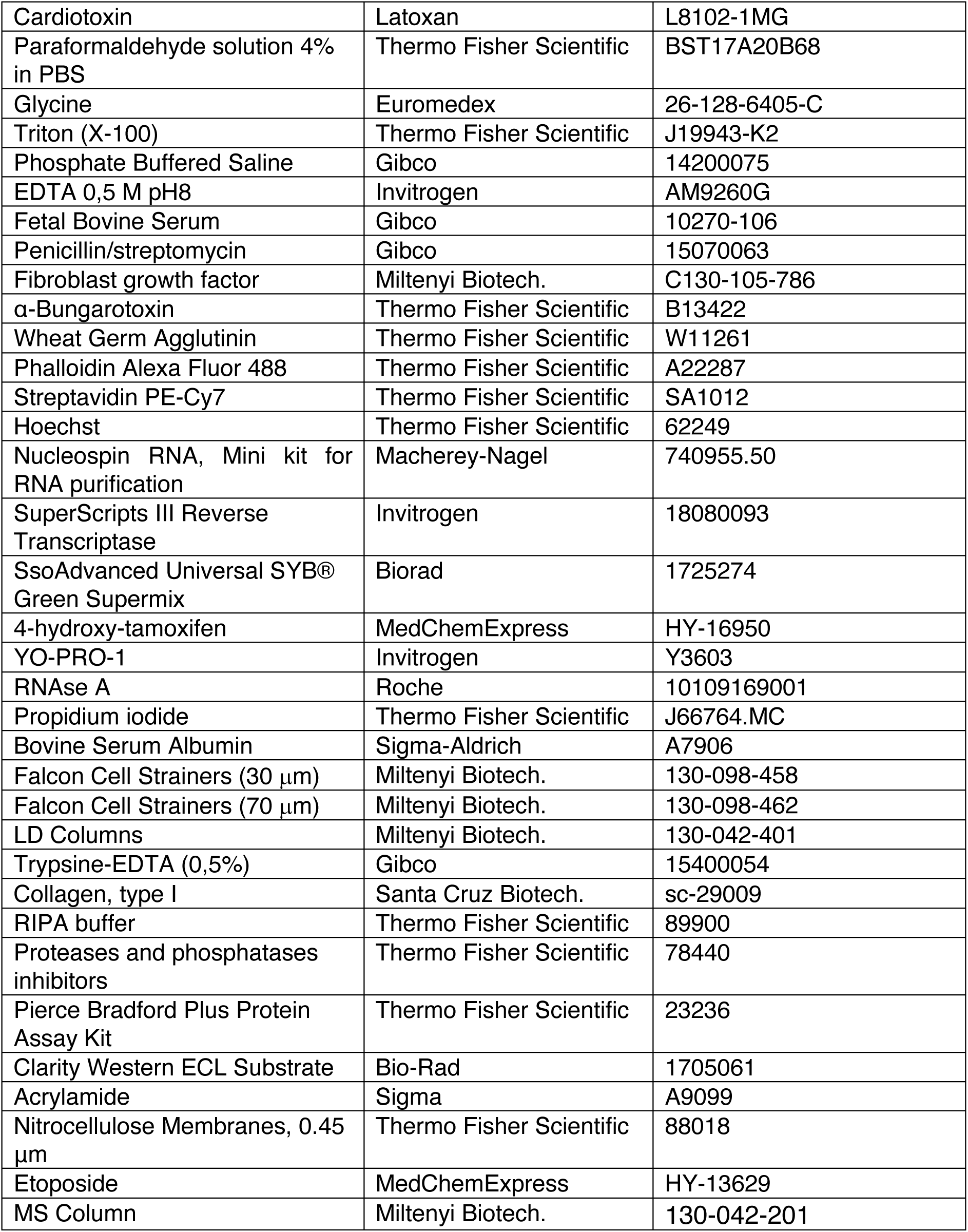

| REAGENT or RESOURCE | SOURCE | IDENTIFIER |
| --- | --- | --- |
| <u>Antibodies</u> |  |  |
| Rat anti-F4/80 | Elabscience | E-AB-F0995A; RRID AB_2539703 |
| Rabbit anti-LAMININα2 | Sigma-Aldrich | L9393; RRID AB_477163 |
| Mouse anti-PAX7 | Santa Cruz | sc-81648; RRID AB_2159836 |
| Goat anti-COLLAGEN TYPE I | Southern Biotech. | 1310-01; RRID AB_2753206 |
| Rat anti-CD31 | Abcam | ab7388; RRID AB_305905 |
| Mouse anti-MyHC (Embryonic) | Santa Cruz Biotech. | sc-53091; RRID AB_670121 |
| Goat Anti-PDGFRα | R&D Systems | AF1062; RRID AB_2236897 |
| Mouse anti-MYOD1 | Santa Cruz Biotech. | sc-377460; RRID AB_2813894 |
| Mouse anti-MyHC (Sarcomeric) | Developmental Studies Hybridoma Bank | MF20; RRID AB_2147781 |
| Anti-CD106 (VCAM1) Biotin clone 429 | Thermo Fisher Scientific | 13-1061-82; RRID AB_466559 |
| Anti-CD45 eFluor450, clone 30-F11 | Thermo Fisher Scientific | 48-0451-80; RRID:AB_1518807 |
| Anti-CD31 eFluor450, clone 390 | Thermo Fisher Scientific | 48-0311-80; RRID:AB_10598808 |
| Anti-CD34 Biotin, clone RAM34 | Thermo Fisher Scientific | 13-0341-81; RRID:AB_466424 |
| Anti-CD64 Alexa Fluor 647, clone X54-5/7.1 | Biolegend | 139322; RRID :AB_2566560 |
| Anti-Ly6A/E(Sca1) Pacific Blue, clone D7 | Biolegend | 108120; RRID:AB_493273 |
| Rabbit anti-Ki67 | Abcam | ab15580; RRID:AB_443209 |
| Rabbit anti-SPARC | Thermo Fisher Scientific | 33-5500; RRID:AB_2533126 |
| Rabbit anti-STING | R&D Systems | MAB7169 RRID:AB_10971940 |
| Rabbit anti-RAB2B | Proteintech | 11756-1-AP; RRID:AB_2176900 |
| Rabbit Anti-cGAS antibody (D1D3G) | Cell Signalling | 15102; RRID:AB_2732795 |
| Rabbit Anti-IRF3 antibody (D83B9) | Cell Signalling | 4302; RRID:AB_1904036 |
| Goat anti-mouse IgG (H+L) cross-adsorbed secondary antibody, Alexa Fluor 647 | Thermo Fisher Scientific | A-21235; RRID:2535804 |
| Goat anti-mouse IgG (H+L) cross-adsorbed secondary antibody, Alexa Fluor 546 | Thermo Fisher Scientific | A-11030;<br>RRID:144695 |
| Goat anti-mouse IgG (H+L) cross-adsorbed secondary antibody, Alexa Fluor 488 | Thermo Fisher Scientific | A-11001;<br>RRID:AB_2534069 |
| Goat anti-mouse IgG (H+L) cross-adsorbed secondary antibody, Alexa Fluor 546 | Thermo Fisher Scientific | A-11010;<br>RRID:AB_2534077 |
| Goat anti-rat IgG (H+L) cross-adsorbed secondary antibody, Alexa Fluor 488 | Thermo Fisher Scientific | A-11006;<br>RRID:AB_2534074 |
| Donkey anti-goat IgG (H+L) cross-adsorbed secondary antibody, Alexa Fluor 488 | Thermo Fisher Scientific | A-11055;<br>RRID:AB_10276611 |
| Goat anti-Mouse IgG (H+L) Secondary Antibody, HRP | Thermo Fisher Scientific | 31430;<br>RRID:AB_228307 |
| Goat anti-Rabbit IgG (H+L) Secondary Antibody, HRP | Thermo Fisher Scientific | 31460;<br>RRID:AB_228341 |
| Deposited data |  |  |
| RNA-Seq Data | GEO | GSE233394 |
| ATAC-seq Data | GEO | GSE233393 |
| Single cell RNA-seq Data | GEO | GSE233395 |
| Mass spectrometry Data | Pride | PXD042002 |
| Differential Gene Expression Analysis and Gene Ontology | Mendeley | https://dx.doi.org/10.17632/r4tj555pc7.1 |
| Experimental models: organisms/strains |  |  |
| B6.Cg-Pax7<sup>tm1(cre/ERT2)Gaka/J</sup> | The Jackson Laboratory | IMSR_JAX:017763 |
| B6.Cg-Tg(Pdgfrα<sup>cre/ERT</sup>)467Dbe/J | The Jackson Laboratory | IMSR_JAX:01828 |
| B6.Setdb1<sup>fox/flox</sup> | Drs. Y. Shinkai, J. Turner | *Matsui et al.*, 2010 |
| Critical commercial assays |  |  |
| Click-iT EdU cell proliferation kit for imaging, Alexa Fluor 488 dye | Invitrogen | C10340 |
| Senescence β-Galactosidase Staining Kit | Cell Signalling | 9860 |
| Satellite cell isolation kit | Miltenyi Biotech | 130-104-268 |
| Anti-Cy5/Anti-Alexa Fluor 647 Microbeads | Miltenyi Biotech | 130-091-395 |
| Chemicals, peptides, and recombinant proteins |  |  |
| Collagenase, type II | Gibco | 17101015 |
| Dispase | Gibco | 17105041 |
| DMEM, low glucose medium | Gibco | 11564446 |
| DMEM, high glucose medium | Gibco | 13345364 |
| Ham's F10 medium | Gibco | 14025092 |
| O.C.T. compound | CellPath | KMA-0100-00A |
| Fluorescent mounting medium | Invitrogen | S3023 |
| Tamoxifen | Sigma-Aldrich | T5648 |
| Cardiotoxin | Latoxan | L8102-1MG |
| Paraformaldehyde solution 4% in PBS | Thermo Fisher Scientific | BST17A20B68 |
| Glycine | Euromedex | 26-128-6405-C |
| Triton (X-100) | Thermo Fisher Scientific | J19943-K2 |
| Phosphate Buffered Saline | Gibco | 14200075 |
| EDTA 0,5 M pH8 | Invitrogen | AM9260G |
| Fetal Bovine Serum | Gibco | 10270-106 |
| Penicillin/streptomycin | Gibco | 15070063 |
| Fibroblast growth factor | Miltenyi Biotech. | C130-105-786 |
| α-Bungarotoxin | Thermo Fisher Scientific | B13422 |
| Wheat Germ Agglutinin | Thermo Fisher Scientific | W11261 |
| Phalloidin Alexa Fluor 488 | Thermo Fisher Scientific | A22287 |
| Streptavidin PE-Cy7 | Thermo Fisher Scientific | SA1012 |
| Hoechst | Thermo Fisher Scientific | 62249 |
| Nucleospin RNA, Mini kit for RNA purification | Macherey-Nagel | 740955.50 |
| SuperScripts III Reverse Transcriptase | Invitrogen | 18080093 |
| SsoAdvanced Universal SYB® Green Supermix | Biorad | 1725274 |
| 4-hydroxy-tamoxifen | MedChemExpress | HY-16950 |
| YO-PRO-1 | Invitrogen | Y3603 |
| RNAse A | Roche | 10109169001 |
| Propidium iodide | Thermo Fisher Scientific | J66764.MC |
| Bovine Serum Albumin | Sigma-Aldrich | A7906 |
| Falcon Cell Strainers (30 μm) | Miltenyi Biotech. | 130-098-458 |
| Falcon Cell Strainers (70 μm) | Miltenyi Biotech. | 130-098-462 |
| LD Columns | Miltenyi Biotech. | 130-042-401 |
| Trypsine-EDTA (0,5%) | Gibco | 15400054 |
| Collagen, type I | Santa Cruz Biotech. | sc-29009 |
| RIPA buffer | Thermo Fisher Scientific | 89900 |
| Proteases and phosphatases inhibitors | Thermo Fisher Scientific | 78440 |
| Pierce Bradford Plus Protein Assay Kit | Thermo Fisher Scientific | 23236 |
| Clarity Western ECL Substrate | Bio-Rad | 1705061 |
| Acrylamide | Sigma | A9099 |
| Nitrocellulose Membranes, 0.45 μm | Thermo Fisher Scientific | 88018 |
| Etoposide | MedChemExpress | HY-13629 |
| MS Column | Miltenyi Biotech. | 130-042-201 |

### RESSOURCE AVAILABILITY

#### Lead contact

Further information and requests for resources and reagents should be directed and will be fulfilled by the lead contact, Fabien Le Grand.

### METHODS DETAILS

#### Mice line

Animals were handled according to the European Community guidelines, implementing the 3Rs rule. Protocols were validated by the ethic committee of the French Ministry, under the reference number APAFIS#34582-202111121631816 v3. All mice were kept under pathogen-free conditions in individually ventilated cages, with a 12h/12h light/dark cycle, and had free access to water and food.

All *in vivo* experiments were performed on 10 weeks old male mice. Generation and genotyping of the *Pax7^CreERT2^*, *Setdb1^lox/lox^* and *Pdgfrα^CreERT2^* lines have been described previously. These strains were used to generate combined experimental strains, which include the *Pax7*^CreERT2/+^:*Setdb1^fl/fl^*, and *PDGFRα^CreERT2^*:*Setdb1^fl/fl^* strains used in the study. Breeding was performed with 2 females and 1 male.

#### Tamoxifen injection and muscle injury

To induce CreER activity, 4 consecutives intraperitoneal (IP) injections of tamoxifen were administrated to the 2 months old mice (100μL at 10mg/mL dissolved in corn oil). Cardiotoxin injection was performed 7 days after the first injection of tamoxifen. Mice were anesthetized with isoflurane inhalation. Acute muscle injury was induced by 50μL intramuscular injections of cardiotoxin (CTX; 10μM) in the *Tibialis Anterior* (TA) muscle.

#### Isolation of TA muscle and Histological analysis

Mice were euthanized with isoflurane asphyxiation followed by cervical dislocation. TA muscles were harvested, weighed, submerged in OCT and then frozen in isopentane cooled in liquid nitrogen. Frozen muscles were stored at −80°C. Muscle cryosections of 10μm thickness were collected using a NX50 cryostat. Fixation of muscle sections was performed with 4% PFA in PBS for 10 minutes at room temperature. Permeabilization was done in 0.1M glycine, 0.1% Triton X-100 diluted in PBS during 10 minutes at room temperature. After 3 washes in PBS, muscle sections were incubated with blocking buffer (3% BSA, 0.1% Triton X-100, in PBS) supplemented with M.O.M. Blocking reagent for 1 hour. Primary antibodies diluted in blocking buffer were incubated overnight at 4°C. Detection of primary antibodies was achieved using Alexa Fluor secondary antibodies diluted at 1:1000 in blocking buffer after 3 washes in PBS. Nuclei were counterstained with Hoechst at 1μg/mL in PBS before mounting with fluorescent mounting medium. Full muscle sections were assembled using a slide scanner ZEISS AxioScan7. Images were processed and analyzed with ZEISS ZEN and FIJI softwares. Myofiber basal lamina was revealed by LAMININα2 immunostaining and cross-sectional analysis was performed using Open-CSAM ImageJ macro^48^. Entire sections were analyzed, giving around 2500 myofibers counted for each analysis.

PAX7+ cells were quantified manually, using the colocalization of nuclear PAX7 immunostaining and Hoechst. COLLAGEN I and F4/80 immunostainings were quantified using FIJI threshold, quantifying the area between the myofibers. Central myonuclei were quantified manually using FIJI cell counter tool.

#### MuSC isolation and cell culture

Skeletal muscle-derived primary myoblasts were isolated from the different mice models, aged between 8- to 16-week-old, using magnetic cell separation (MACS). Hindlimbs muscles from both legs were dissected and placed in a sterile Petri dish containing PBS. Mechanic dissection was performed with scissors to have a pulp of muscle. The pulp was digested in a bath of Collagenase II (2 μg/mL) solution diluted in Ham’s F10 Nutrient Mix during 1 hour at 37°C. A second digestion is performed during 30 min containing Collagenase II and Dispase (3.25μg/mL) at 37°C. The pulp is then filtered (70μm and 30μm) followed by a step of trituration using syringe 0.2G. Washes were done with 5% Bovine Serum Albumin (BSA), 3mM EDTA diluted in phosphate buffered saline (PBS) and myoblast isolation was performed using Satellite Cell Isolation kit (Miltenyi). The cell suspension is then poured into columns placed in a magnetic field (LD Column, Miltenyi). Unlabeled cells (MuSCs) flow through the column while magnetically labeled cells are retained within the column. MuSCs were resuspended in the growth medium (Ham’s F10 with 20% FBS, 1% Penicillin/Streptomycin (P/S)), and 2.5 ng/mL of basic Fibroblast Growth Factor and plated into a collagen-coated 60mm Petri dish. Cells were maintained in the growth medium until cells reached 80% confluence.

Alternatively, after muscle dissection and digestion, cell suspensions were incubated 30 min on ice with the following antibodies: anti-ITGA7, anti-CD34, anti-CD45, Anti-CD106 and anti-Ly-6/E (SCA1). MuSCs were FACS-sorted by gating CD34+/ITGA7+ double positive and CD45-/Ly-6/E-double negative cells.

#### Immunostaining on cell and myofibers

Cell fixation was performed with 4% PFA in PBS for 10 minutes at room temperature and 3 washes in PBS were done before permeabilization step. Permeabilization was done in 0.1M glycine, 0.1% Triton X-100 diluted in PBS for 10 minutes at room temperature. If cells or myofibers were pre-incubated with EdU, the Click-iT reaction was done using the Click-iT EdU Alexa Fluor 647 Imaging kit according to the manufacturer’s protocol. After 3 washes in PBS, muscle section was incubated with blocking buffer for 1 hour. Primary antibodies diluted in blocking buffer were incubated overnight at 4°C. Detection of primary antibodies was achieved using Alexa Fluor secondary antibody at a 1:1000 in blocking buffer after 3 times washes in PBS. Nuclei were counterstained with Hoechst at 1μg/mL in PBS.

#### RNA extraction, cDNA synthesis and RT-qPCR

A minimum of 2 x 10^5^ cells were collected per sample for RNA extraction using the Nucleospin RNA II kit following the manufacturer protocol. RNA concentration was evaluated with Nanodrop. Complementary DNA (cDNA) was generated using High-Capacity Reverse Transcription Kit (Thermo Fisher Scientific). cDNA was then used for quantitative PCR (qPCR) done with SYBR Green Master Mix (Roche) and run in LightCycler 480 for 40 cycles. Primers are reported in Supplementary Table 5. All samples were duplicated, and transcripts levels were normalized for a housekeeping gene relative abundance. The relative mRNA levels were calculated using the 2^−ΔΔCt^ method. The ΔCt were obtained from Ct normalized to the housekeeping genes *Gapdh* or *Hprt* levels in each sample. The Ct between 18 and 28 were considered.

#### Myoblast Differentiation

For differentiation assay, myoblasts were split and seeded at 25,000 cells per cm2 onto Matrigel-coated dishes. The next day, growth medium was replaced by differentiation medium (DMEM with 2% FBS, 1% P/S). Cells were differentiated for 3 days, before fixation and immunostaining. Quantification of muscle nuclei was performed manually in a blinded manner using the point toll counter in ImageJ. The fusion index was calculated as the fraction of nuclei contained within MyHC+ myotubes, which had two or more nuclei, as compared to the number of total nuclei within each ×20 image. The differentiation index was calculated as the fraction of nuclei contained within all MyHC+ cells, including both mononuclear and multinuclear cells, as compared with the number of total nuclei within each ×20 image.

#### Cell death experiment

Myoblasts were cultured in 100mm Petri dish and then treated with either EtOH or 4-OHT for 5 days. Cells were trypsinized and resuspended in PBS. 50μg/mL of PI was added during 30 min at 37°C. After 2 washes, 1μM/mL of YO-PRO-1 was added in PBS during 1 hour on ice. After 2 washes in PBS, cells were analyzed with a flow cytometer BD FACSCanto™ II. The different cell populations were identified using the following gates: the population in the PI-Yo-PRO-1 gate represented the living cells^50^. The population considered as PI+ YO-PRO-1- represented necrotic cells. The population considered as PI-YO-PRO-1+ represented apoptotic cells and PI+ YO-PRO-1+ represented dead cells.

#### Cell cycle experiment using flow cytometry

Myoblasts were cultured in 100mm Petri dish and then treated with either EtOH or 4-OHT for 5 days. At least 1×10^5^ cells were fixed with cold ethanol 70% /PBS. Cells were centrifuged and the pellet was resuspended in 1mL of PBS containing 0.1mg/mL of RNAse A and 50μg/mL of PI overnight at 4°C. After 2 washes in PBS, cells were analyzed with a flow cytometer BD FACSCanto™ II.

#### Western blot

Whole protein extracts were obtained from primary myoblasts using RIPA buffer. The buffer was supplemented with Protease and Phosphatase Inhibitor Cocktail (100X). Protein concentration was determined by Bradford assay. Protein extracts (20 μg) were subjected to SDS–PAGE in 12% polyacrylamide gels, transferred to nitrocellulose membranes, and probed with primary antibodies. Secondary antibodies used were HRP-coupled secondary antibodies (Invitrogen) in a dilution of 1:10000 and Clarity Western ECL Substrate was used to visualize the signal. Western blot densitometry quantification was performed using FIJI software. Briefly, minimum brightness thresholds were increased to remove background signal. Remaining bands were bracketed, plot profiles generated, and area under histograms auto traced. Protein levels were normalized against the levels of the loading control.

#### cGAS inhibitor experiments

RU521 was used at a concentration of 200 ng/ml in the medium and changed every day. Cells *in vitro* were treated at the same time with 4-OHT and RU521 or EtOH and DMSO for 5 days.

#### Macrophages isolation

Cells were extracted from regenerating TA muscle at 4 d.p.i. using the same procedure as for uninjured tissues. Cells were incubated with anti-CD64-AF647 antibody, and then with Anti-Alexa Fluor 647 Microbeads. Following two washes, the cell suspension is then poured into columns placed in a magnetic field (MS Column, Miltenyi). Macrophages retained in the columns are then eluted. Sorted macrophages were cytospun onto glass slides for 5 min at 1500 rpm.

#### Neuro-muscular junction (NMJ) analysis

*Extensor Digitorum Longus* (EDL) single myofiber were isolated as previously published.^49^ EDL were harvested from tendon to tendon and digested for 1 hour in DMEM containing 0.25% collagenase II. Isolated myofibers were fixed using PFA 4% diluted in PBS and stained with Alexa-488-labelled α-Bungarotoxin. The NMJ analysis was done using the *Osseni et al*. method. ^51^

#### SA-βgal experiment

Experiment was performed following the manufacturer’s protocol from the Senescence β-Galactosidase Staining Kit. Positive control was human BJ foreskin fibroblasts cells stimulated with etoposide (12.5 μM, 24 hr) and allowed to recover for 2 days.

#### RNA-sequencing of MuSCs

RNA was prepared as described for RNA extraction and sent to the Genom’IC facility (Institut Cochin, Paris, France). Libraries were prepared with TruSeq Stranded Total Library preparation kit according to supplier recommendations. The important steps of this protocol are the removal of ribosomal RNA fraction from 400ng of total RNA using the Ribo-Zero Gold Kit; fragmentation using divalent cations under elevated temperature to obtain ~300 bp pieces; double strand cDNA synthesis using reverse transcriptase and random primers, and finally Illumina adapters ligation and cDNA library amplification by PCR for sequencing. Sequencing was carried out on pair-end 75 bp of Illumina NextSeq® 500.

#### RNA-sequencing analysis

Nf-core/rnaseq v3.2 was used on a conda container, the raw fastq reads were aligned with STAR and the alignments onto the transcriptome and to perform the downstream BAM-level quantification with Salmon. Genome mapping was done using the genome assembly GRCm38. Data were filtered to conserve only features in canonical chromosomes (1 to 19, X, Y and mitochondrial) identified as protein coding in Ensembl databases. Differential analysis was performed with DEseq2 package (v1.34.0). Gene set enrichment analysis was computed by fgsea package (v1.20.0) on genes ranked by the sign of Log2 Fold Change * log10(p-adjusted by BH). The gProfiler2 package (v.0.2.1) was used to compute and compare multiple input collections from MH, M2:CP, KEGG, M5:GO on genes ranked by the sign of Log2 Fold Change * log10(p-adjusted by BH).

#### ATAC-sequencing of MuSCs

MuSCs were extracted as described in the “myoblast isolation and cell culture”. MuSCs were plated at 10 000 cells/cm^2^ and treated during 3 days with either EtOH or 4-OHT. Cells were trypsinized, frozen and sent to the Diagenode facility (Liège, Belgium) for ATAC and sequencing.

#### ATAC-sequencing analysis

Nf-core/atacseq v1.2.1 was used on a docker container and in standard parameters to compute a rds archive readable in R and different files in multiple formats (bigwig, bed, bam). Data were filtered to conserve only features in canonical chromosomes (1 to 19, X, Y and mitochondrial). To determine chromatin occupancy, bam files were parsed and only reads paired were conserved, properly paired and MAPQ score = 60 with Rsamtools package (v2.10.0). Differential analysis was performed on count reads over regions from peak calling with DEseq2 package (v1.34.0). To facilitate reading, coverage by chromosome was computed, log-transformed coverage scores and plotted only value>3.1 into a circos plot with circlize package (v.0.4.15).

#### Mass spectrometry-based proteomics

MuSCs were treated with EtOH or 4-OHT as described above for ATAC-seq. Cells were thawed and then lysed by RIPA buffer (50 mM Tris-HCl pH7.4, 150 mM sodium chloride, 1 mM EDTA, 0.1% sodium dodecyl sulfate, 1% Triton X-100, 1.5% sodium deoxycholate). Proteins in cell lysates were precipitated by 4 volumes of pre-cooled acetone (Honeywell). The precipitated proteins were sent to the Proteomics core of Biosciences Central Research Facility, HKUST (Clear Water Bay, Hong Kong, China), for subsequent processing. The protein pellets were washed sequentially with pre-cooled acetone, pre-cooled ethanol, and pre-cooled acetone. Sample preparation for mass spectrometry, including protein treatments and clean-up, was done using the iST PreON 96x kit (PreOmics). The samples were loaded onto Bruker timsTOF Pro mass spectrometer. A detailed description of the mass spectrometer is given in ref^53^. The operation parameters had been listed previously^52^. The accumulation and ramp time were set at 100 ms each. The mass spectra were recorded using positive electrospray mode, in the m/z range of 100-1700. The ion mobility was scanned from 0.85-1.35 Vs cm-2. The overall acquisition cycle was 0.53 s, comprising one full TIMS-MS scan and four Parallel Accumulation-Serial Fragmentation (PASEF) MS/MS scans. The TIMS dimension was calibrated linearly using three selected ions from the Agilent ESI LC/MS tuning mix [m/z, 1/K0: (622.0289, 0.9848 Vs.cm^-2^), (922.0097, 1.1895 Vs.cm^-2^), (1221,9906, 1.3820 Vs.cm^-2^)] in positive mode.

#### Mass spectrometry analysis

The raw data were processed by PEAKS software (version: Xpro). The database for protein search was Uniprot. The taxonomy for protein search was Mus musculus. The parent ion tolerance was 15 ppm, and the fragment ion tolerance was 0.06 Da. The protein FDR was 1%. For label-free quantification of the mass spectrometry data, normalized spectral abundance factor (NSAF)-based spectral counting method^54^ was used here. The NSAF method can account for the influence of protein sizes on spectral counts and normalize the total spectral counts across multiple samples. Briefly, spectral abundance factor (SAF) of each protein was obtained by dividing the spectral count with its protein length. Then, NSAF was calculated by dividing the SAF with the total SAF of each sample. Since NSAF values were small, the normalized data were multiplied and presented as NSAF*1000000. To examine differential expression of proteins between control and knockout samples, unpaired two-tailed Student’s t-test was used, assuming equal variance of the two groups. Statistical significance was set at unadjusted p < 0.05.

#### Sample preparation for Single-cell RNA-sequencing

Mice were injected with either corn oil or Tamoxifen and CTX injections on TA were done 7 days after the last injection. TA muscles were harvested 4 days after injury. Cells were extracted from regenerating muscles using the same protocol as for MuSC isolation. Cell suspensions were stained with Hoechst and PI. Live, nucleated cells were FACS sorted and resuspended in encapsulation buffer. Single cell encapsulation and sequencing was performed at the Plateforme Génomique des Cancers facility (Lyon, France) using the 10X Genomic microfluidics system.

#### Single-cell RNA sequencing

Data demultiplexed in fastq format was checked with fastqc and fastq_screen to control quality sequencing. Sequencing reads were processed with STAR (v.2.7.8a) to align and count data for each sample with conservation of rRNA during process. We conserve rRNA data in count matrix. To remove ambient RNA as potential contamination of droplet, SoupX (v.1.6.2) was used on each dataset and new count matrix were built. The downstream analysis was carried to Seurat package (v.4.0.1). Each of the 4 datasets was first analyzed independently before combining and, or resampling datasets. Features in canonical chromosomes (1 to 19, X, Y and mitochondrial), identified as protein coding, mRNA and from mitochondrial in Ensembl databases and expressed at minimum in 10 “cells” were conserved. To compare with others resources, different empiric limitedness from publications used to analyze data from similar single cell experiments on the same whole tissue (800 – 10k counts RNA; 600 – 6000 features RNA; 5% Mitochondrial part). A model of integrity cells was performed to conserve only cells sharing the highest similarity based on a mahalanobis distance on nCount_RNA, nFeature_RNA, percentages of protein coding, mRNA and mitochondrial. Cells with a too high distance were removed. DropletUtils (v.1.14.2) to compute barcode rank statistics and identify knee and inflexion points on the total count curve. Cells were selected when their ranks are between high “empiric” thresholds (10k) and knee plot value for each sample, others cell ranks are rejected. Different categories of cells were calculated based on elbow point on the curve nCount_RNA / nFeature_RNA: low, medium (medium- to medium++) and high. These categories are defined about knee, inflexion, min and max elbow point. Low cell category was rejected. scDblFinder (v.1.80) used to determine singlet to doublet. After filtering, metrics were recalculated.

For KO sample, there are 10,649 singlet detected with [1412:62851] nCount_RNA (median is 6494); [920:7264] nFeature_RNA (median is 2079); [48.85:99.56] %Protein Coding (median is 95.75); [0:50.87] %rRNA (median is 3.68); [0:14.55] %MT (median is 0.44).

For WT sample, there are 8,169 singlet detected with [548:53519] nCount_RNA (median is 9376); [338:7121] nFeature_RNA (median is 2463); [35.10:99.59] %Protein Coding (median is 96.39); [0:64.01] %rRNA (median is 2.97); [0:18.30] %MT (median is 0.51).

A randomized selection and resampling allowed to analyze the same numbers of cells in each sample.

For each sample, SCTransform with glmGamPoi method was done. Variables Feature were selected on a merge sampling and RunPCA on a top3000 features between each sample from Seurat package performed to allow the of DoubletFinder (v.2.0.3) and to identify and to remove cell doublets. The optimal dimension threshold from PCA results were determined by a custom method. Integration was performed on SCT assay with Harmony. We followed the guidelines from Harmony integration.

Before integration, each sample was log-normalized on top32000 features by the vst algorithm. After identifying anchors and performing integration, data were scaled using a linear model. After integration, RunPCA and RunHarmony from the Seurat package were calculated; for both, the optimal dimension threshold was determined by a custom method. The optimal dimension threshold from PCA was used to calculate Harmony reduction. The optimal dimension threshold from Harmony was used to compute the Shared Nearest-neighbor graph.

Original Louvain algorithm with 0.7 resolution was performed to determine clusters. All harmony dimensions were used to Rreduction dimension graphs. Theses graphs were generated by RunTSNE which uses FFT-accelerated Interpolation-based t-SNE (Flt-SNE).

Original Louvain algorithm with 0.7 resolution was performed to determine clusters. Markers of gene expression for each cluster were found by MAST and ROC analysis methods for all features and with no filters (min.pct=0; logfc.threshold=-Inf; only.pos=F; densify=T). Slingshot (v.2.2.1) was used to calculate trajectories. Imbalancescore function was used to compute an imbalance score to show whether nearby cells have the same condition of not and progressionTest function from Condiment package for both (v.0.99.14) was used to compute a test whether or not the pseudotime distribution are identical within lineages between conditions. TradeSeq package (v.1.8.0) was used to allow analysis of gene expression along trajectories. Ucell (v1.2.2) used to calculate score from hallmark mouse genelist.

#### ChIP-sequencing of MuSCs

MuSCs were extracted as described in the “myoblast isolation and cell culture”. MuSCs were plated at 10 000 cells/cm^2^ and treated during 3 days with either ethanol or 4-hydroxytamoxifen. Native ChIP for H3K9me3 and library preparation were performed as previously published. ^6^ Libraries were sequenced on an Illumina Nextseq 500 in a paired-end mode in two independent biological replicates at the Genom’IC facility (Institut Cochin, Paris, France).

#### ChIP-sequencing analysis

Nf-core/chipseq v2.0.0 was used on a singularity container, in standard and custom parameters to compute a rds archive readable in R and different files in multiple formats (bigwig, bed, bam, broadpeak). BWA aligner v0.7.17-r1188 was used, the read length used to calculate MACS2 genome size for peak calling was 50bp and the broad cutoff was 0.1. Broadpeak output files were used to identify broad regions specific to one condition or both. Regions were annotated with ChIPseeker (v1.30.3) from TxDB.Mmusculus.UCSC.mm10.knownGene : the region range of TSS was [-2500;2500]. To determine the nearest repeat element to regions, distances were calculated by distanceToNearest function from GenomicRanges, references used from UCSC rmsk database. Visualization of regions with broadpeak in control-condition and absent broadpeak in KO-condition were managed by Gviz package (v1.38.4). Coverage by chromosome was computed, log-transformed coverage scores into a circos plot with circlize package (v0.4.15).

#### Statistical analysis

A minimum of three biological replicates was performed for the presented experiments. Error bars are standard errors. Statistical significance was assessed by the Student t test, using Microsoft® Excel® and GraphPad Prism 9. Differences were considered statistically significant at the P < 0.05 level. For each sample, four images were taken with a ×4, ×10, or ×20 magnification depending on the experimental design. Cell and western blot quantification and analysis were performed using ImageJ.

## REFERENCES

1. Zakrzewski, W., Dobrzyński, M., Szymonowicz, M., and Rybak, Z. (2019). Stem cells: past, present, and future. Stem Cell Res Ther 10, 68. 10.1186/s13287-019-1165-5.

2. Janssen, S.M., and Lorincz, M.C. (2022). Interplay between chromatin marks in development and disease. Nat Rev Genet 23, 137–153. 10.1038/s41576-021-00416-x.

3. Schultz, D.C., Ayyanathan, K., Negorev, D., Maul, G.G., and Rauscher, F.J. (2002). SETDB1: a novel KAP-1-associated histone H3, lysine 9-specific methyltransferase that contributes to HP1-mediated silencing of euchromatic genes by KRAB zinc-finger proteins. Genes Dev 16, 919–932. 10.1101/gad.973302.

4. Bilodeau, S., Kagey, M.H., Frampton, G.M., Rahl, P.B., and Young, R.A. (2009). SetDB1 contributes to repression of genes encoding developmental regulators and maintenance of ES cell state. Genes Dev 23, 2484–2489. 10.1101/gad.1837309.

5. Ceol, C.J., Houvras, Y., Jane-Valbuena, J., Bilodeau, S., Orlando, D.A., Battisti, V., Fritsch, L., Lin, W.M., Hollmann, T.J., Ferré, F., et al. (2011). The histone methyltransferase SETDB1 is recurrently amplified in melanoma and accelerates its onset. Nature 471, 513–517. 10.1038/nature09806.

6. Zakharova, V.V., Magnitov, M.D., Del Maestro, L., Ulianov, S.V., Glentis, A., Uyanik, B., Williart, A., Karpukhina, A., Demidov, O., Joliot, V., et al. (2022). SETDB1 fuels the lung cancer phenotype by modulating epigenome, 3D genome organization and chromatin mechanical properties. Nucleic Acids Research 50, 4389–4413. 10.1093/nar/gkac234.

7. Yin, H., Price, F., and Rudnicki, M.A. (2013). Satellite cells and the muscle stem cell niche. Physiol. Rev. 93, 23–67. 10.1152/physrev.00043.2011.

8. Porpiglia, E., and Blau, H.M. (2022). Plasticity of muscle stem cells in homeostasis and aging. Current Opinion in Genetics & Development 77, 101999. 10.1016/j.gde.2022.101999.

9. Beyer, S., Pontis, J., Schirwis, E., Battisti, V., Rudolf, A., Le Grand, F., and Ait-Si-Ali, S. (2016). Canonical Wnt signalling regulates nuclear export of Setdb1 during skeletal muscle terminal differentiation. Cell Discovery 2, 16037.

10. Matsui, T., Leung, D., Miyashita, H., Maksakova, I.A., Miyachi, H., Kimura, H., Tachibana, M., Lorincz, M.C., and Shinkai, Y. (2010). Proviral silencing in embryonic stem cells requires the histone methyltransferase ESET. Nature 464, 927–931. 10.1038/nature08858.

11. Murphy, M.M., Lawson, J.A., Mathew, S.J., Hutcheson, D.A., and Kardon, G. (2011). Satellite cells, connective tissue fibroblasts and their interactions are crucial for muscle regeneration. Development 138, 3625–3637. 10.1242/dev.064162.

12. Fasching, L., Kapopoulou, A., Sachdeva, R., Petri, R., Jönsson, M.E., Männe, C., Turelli, P., Jern, P., Cammas, F., Trono, D., et al. (2015). TRIM28 Represses Transcription of Endogenous Retroviruses in Neural Progenitor Cells. Cell Reports 10, 20–28. 10.1016/j.celrep.2014.12.004.

13. Chakkalakal, J.V., Jones, K.M., Basson, M.A., and Brack, A.S. (2012). The aged niche disrupts muscle stem cell quiescence. Nature 490, 355–360. 10.1038/nature11438.

14. Chang, C.C., Chuang, S.-T., Lee, C.Y., and Wei, J.W. (1972). Role of cardiotoxin and phospholipase A in the blockade of nerve conduction and depolarization of skeletal muscle induced by cobra venom. British Journal of Pharmacology 44, 752–764. 10.1111/j.1476-5381.1972.tb07313.x.

15. Cooper, A.J.L., and Hanigan, M.H. (2018). Metabolism of Glutathione S-Conjugates: Multiple Pathways. In Comprehensive Toxicology (Elsevier), pp. 363–406. 10.1016/B978-0-12-801238-3.01973-5.

16. Ramsay, E.E., and Dilda, P.J. (2014). Glutathione S-conjugates as prodrugs to target drug-resistant tumors. Front. Pharmacol. 5. 10.3389/fphar.2014.00181.

17. Karimi, M.M., Goyal, P., Maksakova, I.A., Bilenky, M., Leung, D., Tang, J.X., Shinkai, Y., Mager, D.L., Jones, S., Hirst, M., et al. (2011). DNA Methylation and SETDB1/H3K9me3 Regulate Predominantly Distinct Sets of Genes, Retroelements, and Chimeric Transcripts in mESCs. Cell Stem Cell 8, 676–687. 10.1016/j.stem.2011.04.004.

18. Zhang, S.-M., Cai, W.L., Liu, X., Thakral, D., Luo, J., Chan, L.H., McGeary, M.K., Song, E., Blenman, K.R.M., Micevic, G., et al. (2021). KDM5B promotes immune evasion by recruiting SETDB1 to silence retroelements. Nature 598, 682–687. 10.1038/s41586-021-03994-2.

19. Griffin, G.K. Epigenetic silencing by SETDB1 suppresses tumour intrinsic immunogenicity. 28.

20. Hopfner, K.-P., and Hornung, V. (2020). Molecular mechanisms and cellular functions of cGAS–STING signalling. Nat Rev Mol Cell Biol 21, 501–521. 10.1038/s41580-020-0244-x.

21. Fajgenbaum, D.C., and June, C.H. (2020). Cytokine Storm. N Engl J Med 383, 2255– 2273. 10.1056/NEJMra2026131.

22. Satija, R., Farrell, J.A., Gennert, D., Schier, A.F., and Regev, A. (2015). Spatial reconstruction of single-cell gene expression data. Nat Biotechnol 33, 495–502. 10.1038/nbt.3192.

23. Rosvall, M., and Bergstrom, C.T. (2011). Multilevel Compression of Random Walks on Networks Reveals Hierarchical Organization in Large Integrated Systems. PLoS ONE 6, e18209. 10.1371/journal.pone.0018209.

24. Street, K., Risso, D., Fletcher, R.B., Das, D., Ngai, J., Yosef, N., Purdom, E., and Dudoit, S. (2018). Slingshot: cell lineage and pseudotime inference for single-cell transcriptomics. BMC Genomics 19, 477. 10.1186/s12864-018-4772-0.

25. Galluzzo, C., Chiapparoli, I., Corrado, A., Cantatore, F.P., Salvarani, C., and Pipitone, N. (2023). Rare forms of inflammatory myopathies - part II, localized forms. Expert Review of Clinical Immunology 19, 185–191. 10.1080/1744666X.2023.2154655.

26. Chung, M.-I., Bujnis, M., Barkauskas, C.E., Kobayashi, Y., and Hogan, B.L.M. (2018). Niche-mediated BMP/SMAD signaling regulates lung alveolar stem cell proliferation and differentiation. Development 145, dev163014. 10.1242/dev.163014.

27. Chuong, E.B., Elde, N.C., and Feschotte, C. (2017). Regulatory activities of transposable elements: from conflicts to benefits. Nat Rev Genet 18, 71–86. 10.1038/nrg.2016.139.

28. De Cecco, M., Criscione, S.W., Peterson, A.L., Neretti, N., Sedivy, J.M., and Kreiling, J.A. (2013). Transposable elements become active and mobile in the genomes of aging mammalian somatic tissues. Aging 5, 867–883. 10.18632/aging.100621.

29. Valdebenito-Maturana, B., Valdebenito-Maturana, F., Carrasco, M., Tapia, J.C., and Maureira, A. (2022). Activation of Transposable Elements in Human Skeletal Muscle Fibers upon Statin Treatment. IJMS 24, 244. 10.3390/ijms24010244.

30. Schoggins, J.W., Wilson, S.J., Panis, M., Murphy, M.Y., Jones, C.T., Bieniasz, P., and Rice, C.M. (2011). A diverse range of gene products are effectors of the type I interferon antiviral response. Nature 472, 481–485. 10.1038/nature09907.

31. Haag, S.M., Gulen, M.F., Reymond, L., Gibelin, A., Abrami, L., Decout, A., Heymann, M., van der Goot, F.G., Turcatti, G., Behrendt, R., et al. (2018). Targeting STING with covalent small-molecule inhibitors. Nature 559, 269–273. 10.1038/s41586-018-0287-8.

32. de Oliveira Mann, C.C., Orzalli, M.H., King, D.S., Kagan, J.C., Lee, A.S.Y., and Kranzusch, P.J. (2019). Modular Architecture of the STING C-Terminal Tail Allows Interferon and NF-κB Signaling Adaptation. Cell Reports 27, 1165–1175.e5. 10.1016/j.celrep.2019.03.098.

33. Glück, S., Guey, B., Gulen, M.F., Wolter, K., Kang, T.-W., Schmacke, N.A., Bridgeman, A., Rehwinkel, J., Zender, L., and Ablasser, A. (2017). Innate immune sensing of cytosolic chromatin fragments through cGAS promotes senescence. Nat Cell Biol 19, 1061– 1070. 10.1038/ncb3586.

34. Gulen, M.F., Koch, U., Haag, S.M., Schuler, F., Apetoh, L., Villunger, A., Radtke, F., and Ablasser, A. (2017). Signalling strength determines proapoptotic functions of STING. Nat Commun 8, 427. 10.1038/s41467-017-00573-w.

35. Zierhut, C., Yamaguchi, N., Paredes, M., Luo, J.-D., Carroll, T., and Funabiki, H. (2019). The Cytoplasmic DNA Sensor cGAS Promotes Mitotic Cell Death. Cell 178, 302–315.e23. 10.1016/j.cell.2019.05.035.

36. Chen, H., Chen, H., Zhang, J., Wang, Y., Simoneau, A., Yang, H., Levine, A.S., Zou, L., Chen, Z., and Lan, L. (2020). cGAS suppresses genomic instability as a decelerator of replication forks. Sci Adv 6, eabb8941. 10.1126/sciadv.abb8941.

37. Theret, M., Mounier, R., and Rossi, F. (2019). The origins and non-canonical functions of macrophages in development and regeneration. Development 146, dev156000. 10.1242/dev.156000.

38. Fu, G., Wu, Y., Zhao, G., Chen, X., Xu, Z., Sun, J., Tian, J., Cheng, Z., Shi, Y., and Jin, B. (2022). Activation of cGAS-STING Signal to Inhibit the Proliferation of Bladder Cancer: The Immune Effect of Cisplatin. Cells 11, 3011. 10.3390/cells11193011.

39. Bao, T., Liu, J., Leng, J., and Cai, L. (2021). The cGAS–STING pathway: more than fighting against viruses and cancer. Cell Biosci 11, 209. 10.1186/s13578-021-00724-z.

40. Tang, C.-H.A., Zundell, J.A., Ranatunga, S., Lin, C., Nefedova, Y., Del Valle, J.R., and Hu, C.-C.A. (2016). Agonist-Mediated Activation of STING Induces Apoptosis in Malignant B Cells. Cancer Res 76, 2137–2152. 10.1158/0008-5472.CAN-15-1885.

41. Ou, L., Zhang, A., Cheng, Y., and Chen, Y. (2021). The cGAS-STING Pathway: A Promising Immunotherapy Target. Front. Immunol. 12, 795048. 10.3389/fimmu.2021.795048.

42. Wang, Q., Bergholz, J.S., Ding, L., Lin, Z., Kabraji, S.K., Hughes, M.E., He, X., Xie, S., Jiang, T., Wang, W., et al. (2022). STING agonism reprograms tumor-associated macrophages and overcomes resistance to PARP inhibition in BRCA1-deficient models of breast cancer. Nat Commun 13, 3022. 10.1038/s41467-022-30568-1.

43. Qian, B.-Z., and Pollard, J.W. (2010). Macrophage Diversity Enhances Tumor Progression and Metastasis. Cell 141, 39–51. 10.1016/j.cell.2010.03.014.

44. Miao, L., Qi, J., Zhao, Q., Wu, Q.-N., Wei, D.-L., Wei, X.-L., Liu, J., Chen, J., Zeng, Z.-A., Ju, H.-Q., et al. (2020). Targeting the STING pathway in tumor-associated macrophages regulates innate immune sensing of gastric cancer cells. Theranostics 10, 498–515. 10.7150/thno.37745.

45. Zhang, R.-H., Judson, R.N., Liu, D.Y., Kast, J., and Rossi, F.M.V. (2016). The lysine methyltransferase Ehmt2/G9a is dispensable for skeletal muscle development and regeneration. Skeletal Muscle 6, 22. 10.1186/s13395-016-0093-7.

46. Biferali, B., Bianconi, V., Perez, D.F., Kronawitter, S.P., Marullo, F., Maggio, R., Santini, T., Polverino, F., Biagioni, S., Summa, V., et al. (2021). Prdm16-mediated H3K9 methylation controls fibro-adipogenic progenitors identity during skeletal muscle repair. Sci. Adv. 7, eabd9371. 10.1126/sciadv.abd9371.

47. Bulut-Karslioglu, A., De La Rosa-Velázquez, I.A., Ramirez, F., Barenboim, M., Onishi-Seebacher, M., Arand, J., Galán, C., Winter, G.E., Engist, B., Gerle, B., et al. (2014). Suv39h-Dependent H3K9me3 Marks Intact Retrotransposons and Silences LINE Elements in Mouse Embryonic Stem Cells. Molecular Cell 55, 277–290. 10.1016/j.molcel.2014.05.029.

48. Desgeorges, T., Liot, S., Lyon, S., Bouvière, J., Kemmel, A., Trignol, A., Rousseau, D., Chapuis, B., Gondin, J., Mounier, R., et al. (2019). Open-CSAM, a new tool for semi-automated analysis of myofiber cross-sectional area in regenerating adult skeletal muscle. Skeletal Muscle 9, 2. 10.1186/s13395-018-0186-6.

49. Brun, C.E., Wang, Y.X., and Rudnicki, M.A. (2018). Single EDL Myofiber Isolation for Analyses of Quiescent and Activated Muscle Stem Cells. In Cellular Quiescence Methods in Molecular Biology., H. D. Lacorazza, ed. (Springer New York), pp. 149–159. 10.1007/978-1-4939-7371-2_11.

50. Allen, P., and Davies, D. (2007). Apoptosis Detection by Flow Cytometry. In Flow Cytometry, M. G. Macey, ed. (Humana Press), pp. 147–163. 10.1007/978-1-59745-451-3_6.

51. Osseni, A., Ravel-Chapuis, A., Thomas, J.-L., Gache, V., Schaeffer, L., and Jasmin, B.J. (2020). HDAC6 regulates microtubule stability and clustering of AChRs at neuromuscular junctions. Journal of Cell Biology 219, e201901099. 10.1083/jcb.201901099.

52. Zeng, W., Yue, L., Lam, K.S.W., Zhang, W., So, W.-K., Tse, E.H.Y., and Cheung, T.H. (2022). CPEB1 directs muscle stem cell activation by reprogramming the translational landscape. Nat Commun 13, 947. 10.1038/s41467-022-28612-1.

53. Meier, F., Brunner, A.-D., Koch, S., Koch, H., Lubeck, M., Krause, M., Goedecke, N., Decker, J., Kosinski, T., Park, M.A., et al. (2018). Online Parallel Accumulation–Serial Fragmentation (PASEF) with a Novel Trapped Ion Mobility Mass Spectrometer. Molecular & Cellular Proteomics 17, 2534–2545. 10.1074/mcp.TIR118.000900.

54. Zhu, W., Smith, J.W., and Huang, C.-M. (2010). Mass Spectrometry-Based Label-Free Quantitative Proteomics. Journal of Biomedicine and Biotechnology 2010, 1–6. 10.1155/2010/840518.

